# Intra- and interspecific variations in flight performance of oak-associated Agrilinae (Coleoptera: Buprestidae) using computerised flight mills

**DOI:** 10.1101/2024.07.01.601558

**Authors:** Elodie Le Souchu, Aurélien Sallé, Stéphanie Bankhead-Dronnet, Mathieu Laparie, Daniel Sauvard

## Abstract

Several Agrilinae species (Coleoptera: Buprestidae) are secondary pests of broadleaf forests, and some of them are also major invasive pests. These thermophilous borers are expected to be favoured by climate change and the global deterioration of forest health, and ultimately expand their range and damage. Flight behaviour and performance of these insects are poorly known despite their critical role in dispersal inside and outside native ranges and their relevance for management purposes. This study aimed to assess intra- and interspecific variability in active flight of several Agrilinae species and effects of sex and mass on this variability to contribute to filling this gap. We assessed the flight performance of eleven species associated with oaks (nine *Agrilus*, one *Coraebus* and one *Meliboeus*) plus one *Agrilus* species associated with the herbaceous layer. Computer-monitored flight mills were used to measure flight parameters, including periods, durations, distances and velocity in 250 beetles. Overall, flight capacities were rather homogeneous among species, with a dominance of poor flyers and only *Coraebus undatus* showed outstanding performance. Beetles generally performed several short flight bouts within one trial, and only a few individuals sustained long flight. The maximal total distance covered across multiple assays until death ranged from 170 to 16 097 m depending on the species, with a median between 35 and 966 m (excluding individuals that never flew). On top of this interspecific variability, flight distances also varied greatly among individuals, but were not influenced by sex. Preflight body mass had mixed effects depending on the species, presumably related to different dispersal patterns. In our experimental conditions, most species had limited average dispersal capacities over multiple flight trials. Overall, dispersal over long distances and colonisation events probably depend mainly on a small proportion of individuals which largely exceeded the median performance.

## Introduction

The jewel beetle subfamily Agrilinae (Coleoptera: Buprestidae) includes numerous species, especially in the genus *Agrilus* which gathers more than 3 300 described species worldwide (Jendek & Grebennikov, 2023). Some of these thermophilous and heliophilous insects (Reed et al., 2018) are known to be phytosanitary threats on a wide diversity of woody and some non-woody plants (Jendek & Poláková, 2014) and can have important economic and ecological impacts in their native area. Larvae develop in the roots, stems, trunks or branches of their host plant. Within trees, larval development could typically last several years. Imaginal moult takes place in spring, within host tissues. After emergence, they usually move towards the foliage to perform a maturation feed and mate. Then females search for a suitable host to lay their eggs (Bílý, 2002).

As opportunistic pests, Agrilinae are associated with forest declines and diebacks (Le Souchu et al., 2024). In temperate forests, wood-borers such as *Agrilus biguttatus* (Fabricius) and *Agrilus viridis* (Linnaeus) are aggravating agents of large scale oak and beech forest declines (Sallé et al., 2014; Brown et al., 2015; Brück-Dyckhoff et al., 2019; Macháčová et al., 2022). A related Agrilinae species, *Coraebus undatus* (Fabricius) has also been associated with oak decline (Du Merle & Attié, 1992; Sallé et al., 2014). *Agrilus* species are also known for their high invasive potential, supported by several examples of human-mediated invasions (e.g., Wu et al., 2017; Hızal & Arslangündoğdu, 2018; Bozorov et al., 2019). Indeed, their long and cryptic pre-imaginal life in the wood facilitates unnoticed transportations and introductions (Chamorro et al., 2015; Ruzzier et al., 2023). One of the prominent invasive pests worldwide is the Emerald Ash Borer (EAB), *Agrilus planipennis* Fairmaire. It is responsible for most of the worldwide economic damage attributed to Buprestidae (Renault et al., 2022). This species has invaded North America from Asia (Herms & McCullough, 2014) and arrived in Russia in 2003, now threatening the western part of Europe (Orlova-Bienkowskaja et al., 2020). Other *Agrilus* species have been identified as potential future invaders, such as *Agrilus anxius* (Evans et al., 2020) or *Agrilus fleischeri* Obenberger (Volkovitsh et al., 2020), including species belonging to the European assemblage of oak-associated borers. Current forest decline phenomena are beneficial to these borers (Le Souchu et al., 2024), together with climate change which is suspected to assist their northward expansion (Pederson & Jørum, 2009; Brown et al., 2015). Within this context, evaluating flight capacities of intercepted or introduced species is of paramount importance to evaluate their dispersal capacities and implement efficient monitoring and control strategies.

Active flight is a central function in the life strategies of most insects (Dickinson et al., 1999; Dudley, 2002). This mode of locomotion is involved in the search for a mate, trophic resources or favourable environmental conditions, as well as in dispersal and colonisation of new geographic areas (Dudley, 2002). The distance covered during the lifetime of an individual can vary considerably among genera, species, and even individuals. It can be influenced by numerous factors, ranging from intrinsic features (physiological traits, reserves, morphological characteristics and behaviours, mated status, *etc.*), to external (a)biotic factors (temperature, crowding, food availability, *etc.*) and their spatiotemporal variations, immediate or not (Minter et al., 2018). The study of flight capacities and their variations at multiple focal scales can provide valuable insights to better understand the ecology of species and, in particular, help to estimate their dispersal potential (see e.g., Robinet et al., 2012). Yet, in the case of *Agrilus* spp., despite their economic and ecological damage and their predisposition to be accidentally introduced into new environments, this information remains scarce in the literature. Their flight capacities have rarely been tested, let alone in European species. Therefore, the numerous works on *A. planipennis* are used to estimate the expected dispersal capacities of potential invasive *Agrilus* spp. or introduced ones (e.g., Baranchikov et al., 2019a; b) and highlight their high flight capacities which may facilitate further dispersal (Taylor et al., 2010; USDA-APHIS, 2011). While male *Agrilus* may be more prone to staying close to their birth host to look for a mate (Rodriguez-Soana et al., 2007), gravid females are able to disperse several kilometres away (Taylor et al., 2010). However, it is often considered that most females will disperse only tens of meters if the resource distribution allows it (Mercader et al., 2009).

Several methods exist to estimate insect flight capacities. Laboratory approaches are usually favoured (Minter et al., 2018; Naranjo, 2019) due to the short lifespan of insects and the difficulty to track them in the field visually or using mark-recapture protocols; in addition, most insects, including *Agrilus* spp., are too tiny to carry even the smallest tags. These approaches use either free flight protocols (or free flight chamber, Taylor et al., 2010; e.g., wind tunnel, Crall et al., 2017) or tethered techniques (e.g., flight mill, Sauvard et al., 2018). The latter are usually preferred, especially flight mills, which are rather inexpensive, produce a wealth of data, have no limit on flying distance, and are easy to replicate under controlled conditions. As insects tethered on a flight mill never reach any target, flight mills provide information on flight capacities rather than actual flight dynamics and behaviours *in natura*. Their advantages are therefore offset by the fact that they create artificial conditions, with multiple potential biases. However, their easy replication allows comparing individuals and species in the exact same conditions, and many authors showed that tethered flight results are consistent with general knowledge on the studied insect performances, or even actual field flight capacity when it is available (Robinet et al., 2012, on processionary moth; Sauvard et al., 2018, on yellow-legged hornet).

In this study, we aimed at determining flight capacities of several *Agrilus* species, and specifically their intra- and interspecific variations. The focus on *Agrilus* behaviour and biology may help establish management strategies and improve detection tools. We measured flight potential of oak-associated species using wild individuals tethered to computer-monitored flight mills. An herbaceous species, *Agrilus hyperici* (Creutzer), similar in size to most of the oak-associated species we sampled, was also added to the pool of species. This species was intentionally introduced in North America from France in 1953 as a biological control agent to target *Hypericum perforatum* L. and its dispersal have been monitored (Jendek & Grebennikov, 2009; Gov. of British Columbia, 2018). This addition allowed to (1) confirm that the performance of *A. hyperici* deals with the knowledge on its dispersion *in natura*, and (2) compare flight performance of forest species to that of an herbaceous-associated species. First, we expected noticeable flight differences among species, as previously observed within multiple coleopteran families (Nitidulidae, Van Dam et al., 2000; Cetoniinae, Dubois et al., 2009). Second, we hypothesised high inter-individual variations in flight measurements, especially a strong sexual dimorphism with females putatively flying greater distances than males, as is the case for mated females of *A. planipennis* (Taylor et al., 2010).

## Material and methods

### Sampling and rearing design

We conducted three consecutive years of experiments, from 2021 to 2023, in France. Live *Agrilus* were caught using two methods. First, we used green multi-funnel traps (ChemTica Internacional, San Jose, Costa Rica) with twelve fluon-coated funnels. These traps are commonly used to sample Agrilinae (Rassati et al., 2019; Le Souchu et al., 2024). From June 23 to August 06 in 2021 and from May 09 to July 27 in 2022, respectively six and eight traps were set at the University of Orléans (Longitude: 47.85 N, Latitude: 1.94 E; WGS 84, DDD), and four and six traps at the INRAE site of Ardon (47.83 N, 1.91 E). These two sites are separated by 2.6 km. Selected forest patches in these areas were dominated by *Quercus robur* (L.) and *Quercus petraea* (Matt.) Liebl. A fresh oak leaf was placed in each collector to provide food and support to trapped insects, and no preservative liquid was used in order to keep them alive. Traps were hung among the lower branches in the canopy (i.e., approximately 15 m above the ground), to capture adults after their emergence, during their search for a mate or a suitable host. No lure was added to the traps and the collectors were emptied every morning during weekdays. Additionally, we captured *A. hyperici* on *H. perforatum* and oak borers on oak logs using a sweep net. In 2022, *A. hyperici* were sampled in three locations, at Saint-Denis-en-Val (47.89 N, 1.98 E) on June 26 and 28, at the University of Orléans on July 04, and at the INRAE site of Ardon on July 06. In 2023, we similarly captured *A. hyperici* in Vouzon (47.64 N, 2.05 E) on June 16. The same day, several specimens of oak borers were captured in a log deposit nearby (47.65 N, 2.01 E). All combined, the oak-associated species caught were: *Agrilus angustulus* (Illiger), *Agrilus graminis* Kiesenwetter, *Agrilus hastulifer* (Ratzeburg), *Agrilus laticornis* (Illiger), *Agrilus sulcicollis* Lacordaire, *C. undatus, Meliboeus fulgidicollis* (P.H. Lucas); and species specialised in others hosts (indicated between parentheses) were: *Agrilus graecus* Obenberger (*Viscum* spp.), *Agrilus olivicolor* Kiesenwetter (*Carpinus* spp.), *A. viridis* (*Fagus* spp.) and *A. hyperici* (*H. perforatum*). All individuals are listed in the supplementary table S1.

Live Agrilinae collected were maintained in the INRAE site of Ardon, in a climatic chamber (Galli GTEST-CL-0700) at a 12:8-hour cycle of 25:20 °C ± 0.2 °C with 2-hour gradients between both phases, 16:8-hour L:D photoperiod using three LED spots (Lédis, 3.5 W, 300 lumens, 4500 K), and 60 % ± 5 % relative humidity (warm phase and photophase were centred on solar midday, 2 pm local time). Each insect was put in an individual plastic vial capped with wet cotton to allow ventilation and provide a water source. Since species were determined only after death, all vials but those containing *A. hyperici* (which were known in advance due to the different sampling method) were supplied with a fresh oak leaf renewed every two days, whereas vials containing *A. hyperici* were supplied with fresh *H. perforatum* leaves.

### Flight mill design

Rotational flight mills were designed to estimate beetle flight capacities. Each flight mill (Figure 1) was built on a wooden square base (19 × 19 cm), with an upwards wooden stand in the centre and a wooden bracket on one corner, extending up to the centre. A magnet (Supermagnete, https://www.supermagnete.de/, S-05-05-N) was placed under the bracket and another one on the stand, both vertically aligned to hold an entomological pin (size 2). To minimise friction, the pinhead had no contact with the lower magnet, and only the point of the pin was in contact with the upper magnet. The pin supported a horizontal 16-cm arm, composed of two identical 1-mm-diameter carbon rods connected by a 2-mm-diameter 2-cm aluminium tube; a hole was drilled in the middle of the tube, through which the pin was inserted and glued. The arm carried a small magnet (Supermagnete, S-03-02-N) glued on the lower side, near one end of the tube. The insect was placed at the end of the arm opposite to this magnet and drove the arm while flying. For heavy species (*C. undatus*), a counterweight made of reusable adhesive paste was also placed on the arm end opposite to the insect.

**Figure 1.**
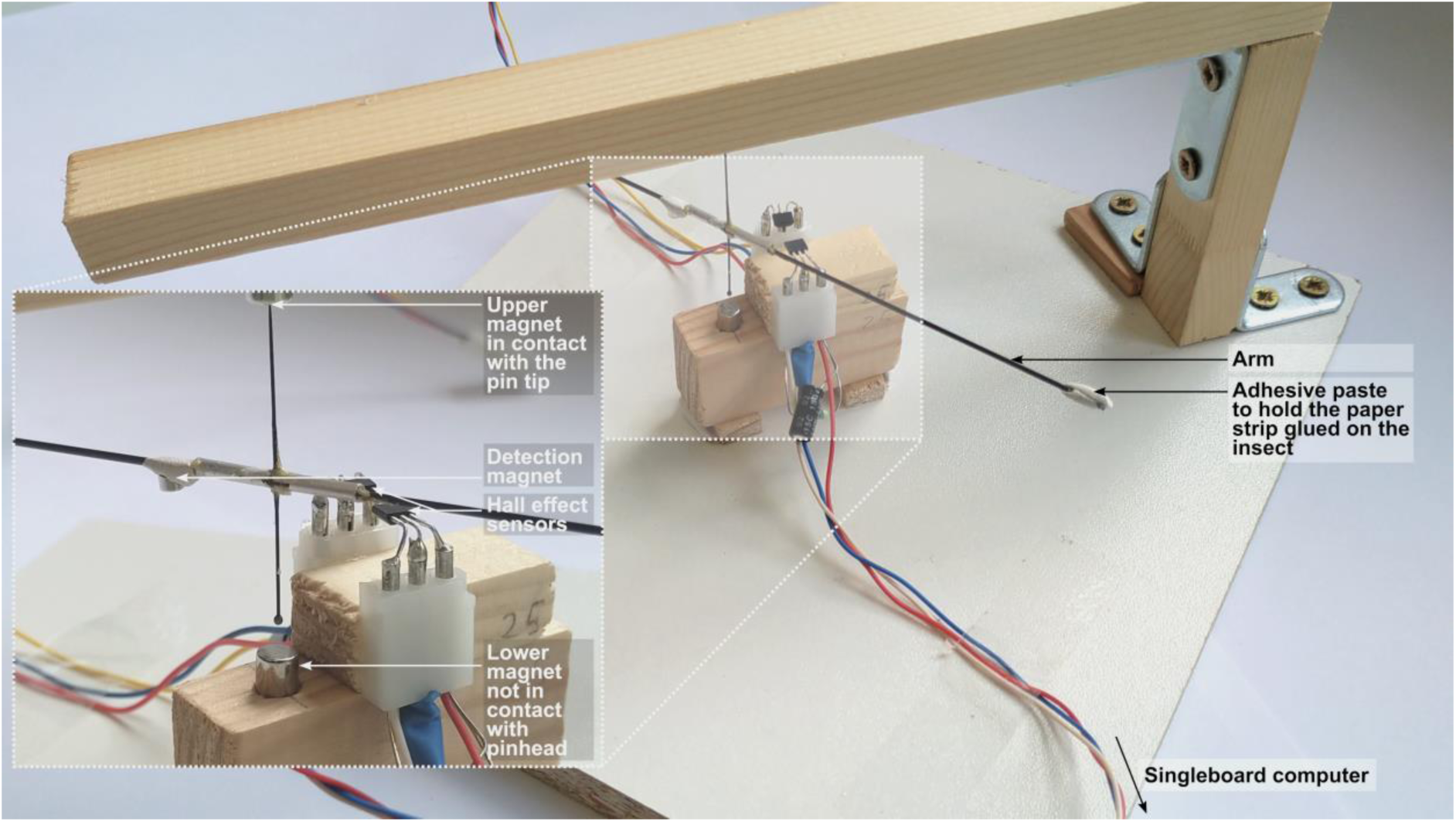
Flight mill used for this study, with a close-up view of the sensor and axis part. The electronics of the system are not shown.

A complete lap corresponded to a travel distance of 0.5 metre. Arm moves were tracked using two Hall effect sensors (DiodesZetex AH337-PG-B) able to detect magnetic fields. They were positioned on the stand just below the arm, approximately 60° from each other around the pin, and detected every small magnet transit over them. Each Hall sensor was connected to a single board computer (Raspberry Pi), to which signals were transmitted. A total of 26 flight mills were built, and managed using 4 Raspberry Pi (1 Raspberry Pi B+, managing 2 flight mills; 1 Raspberry Pi 2 B, 1 Raspberry Pi 3 B and 1 Raspberry Pi 3 B+, each managing 8 flight mills).

An in-house Crystal program running on these computers managed the signals. The program initialised each flight test with flight user-set metadata (mill identifier, insect identifier and gender if known in advance), and start time, creating two dedicated CSV files (one per Hall sensor), hereafter referred to as log files. Then, each associated signal detection was time-stamped and recorded in the appropriate file. At the end of the test, end time was also recorded and the log files were closed.

Flight mills were installed in climate chambers (three PHCBi MIR-254-PE, with six flight mills each, and two Galli GTEST-TM-0700, with four flight mills each; 25 °C ± 0.5 °C, 50 % ± 10 % relative humidity). Each chamber shelf carried a pair of flight mills, which were lit by two LED spots (same as in the rearing chamber). In order to stimulate flight, fresh oak branches with leaves (or *H. perforatum* branches when *A. hyperici* was tested) were placed on each shelf, between the flight mills.

### Experimental design

As Agrilinae are diurnal and thermophilous insects (Reed et al., 2018), we tested them for flight capacities during daytime and at 25 °C ± 0.5 °C. After its sampling, each beetle was maintained one day in rearing conditions (initial rest). The first flight test occurred the next working day; it lasted eight hours. The beetle was then tested again on the flight mill each working day until its death, but these flight tests lasted only four hours. However, test frequency was lowered to a test per week after the tenth one. Test time was chosen so that insects were extracted from the rearing chamber at least an hour after the beginning of the warm phase of their rearing thermal cycle; first tests started at approximately 10 am, and next tests at about 2 pm. In 2021, due to initial low availability of flight mills, only the first test was done with some beetles; as two of them remained alive three weeks after, they had then been tested (during 4 hours) each working day until their death.

Beetles were weighed on a microbalance (Mettler Toledo AG204, d = 0.1 mg) before each flight test. They were then placed in a 3 °C walk-in chamber to slow them down and ease glueing a strip of paper narrower than their width on the dorsal side of their pronotum using cyanoacrylate glue, the most common glue used to tether insects (Naranjo, 2019). The length and width of strips were adjusted to the dimensions of insects. This operation in cold conditions lasted from 15 to 30 minutes, then insects were returned to ambient temperature and immediately attached to flight mills with reusable adhesive paste between the mill’s arm and the strip of paper. This setup allowed rapidly installing and removing tested insects at the beginning and the end of experiments. At the end of flight tests, paper strips were delicately removed from insects, as well as any visible residue of glue, and insects were weighed again before being returned to their rearing vial and chamber. However, no clear variation of the fresh mass was observed during a flight mill trial or during beetle life and only the initial preflight mass was used.

### Data analysis

In-house *Ruby* scripts were developed to analyse the log files generated by the *Crystal* program used to monitor flight mills. They gathered data from the two log files generated by a flight mill during a test, separated flight phases (identified using unequal alternance of signals from the two Hall sensors) from noise and oscillations, and recorded their timestamp, duration and length in the raw dataset used in further analyses. The flight phases were hereafter referred to as flight bouts.

We focused our study on several measures of flight capacities, traditionally used to assess flight performance, which included total flight distance (m), mean flight bout distance (m), total flight duration (s), mean flight bout duration (s), number of flight bouts and latency to first flight bout (s). The latency to first flight bout was used as a measure of the propensity to fly. However, several measures were correlated and we chose to not include the mean flight bout duration and the total flight duration in analyses. The selection method with the correlation tests are available as supplementary material (Document S2). We also explored the relationship between the fresh body mass before the first flight mill trial, hereafter referred to as preflight mass, with the performance. For statistical reasons, species with fewer than six individuals in total were removed from all per-species analyses or figures, and only included in approaches where all species were pooled together. Some flight tests experienced technical problems, such as insect escape or electronic issues, and were excluded from analyses. As indicated above, test frequency changed after the tenth test, and, in 2021, two beetles completed their tests, except the first one, after a long rest period; the corresponding data, hereafter referred to as unusual trial frequency data, was also often excluded from analyses.

Standard linear model (LM) was used to assess linear regression. Kruskal-Wallis tests were used to assess the difference in flight capacities between groups. The Benjamini-Hochberg method permitted to correct p-values for False Discovery Rate. Kruskal-Wallis tests were followed by a post-hoc Dunn test with the Holm adjustment method.

LMERs were used to evaluate the effect of sex and preflight body mass on response variables associated with flight performance of the three most abundant species (*A. hyperici, A. laticornis* and *A. olivicolor*). Every response variable was log-transformed to improve the models, and Q-Q plots were used to review them. Since data from all three years were analysed together, the year of experiment (i.e., 2020, 2021, 2022) was included as a random effect. For each response variable, the null model was compared to the model with preflight mass and sex as explanatory variables, based on their respective AICc (Akaike Information Criterion scores) with a correction for small sample sizes.

When it was needed to compare data from the first eight-hour trial with the subsequent four-hour trials, we truncated the former one to four hours. This resulted in only limited data discarded, as on average 83 % of total distance of eight-hour trials is flown during the first four hours. Unusual trial frequency data was discarded from analyses.

All analyses and figures were realised using *R* 4.2 and *R* 4.3 (R Core Team, 2022, 2023).

## Results

### General beetle characteristics

A total of 250 beetles were tested during the three-year experiment, with 102 individuals in 2021, 125 in 2022, and 23 in 2023 (Table 1). Twelve species were collected, among which three dominated the samples: *A. laticornis*, *A. hyperici* and *A. olivicolor*. Sex ratio was found to be relatively balanced in the three most numerous species, and often biased towards fewer males than females in rarer species, but their small sample sizes warrants caution on this result.

**Table 1.** Sample sizes for each species and sex over the three-year experiment. Values outside brackets indicate the number of individuals for each species and sex collected and tested in flight mills, and values within brackets indicate the number of those insects that could be incorporated in analyses and figures for the first trial. The four rarest species (indicated by an asterisk) were discarded from all analyses or figures that included a species factor.

|  | 2021 |  |  |  | 2022 |  |  |  | 2023 |  |  |  | Total |  |
| --- | --- | --- | --- | --- | --- | --- | --- | --- | --- | --- | --- | --- | --- | --- |
| Species | Females |  | Males |  | Females |  | Males |  | Females |  | Males |  |  |  |
| *Agrilus angustulus* | 1 | (0) |  |  | 1 | (1) | 2 | (2) | 5 | (5) | 2 | (2) | 11 | (10) |
| *Agrilus graecus* * |  |  | 1 | (0) |  |  |  |  |  |  |  |  | 1 | (0) |
| *Agrilus graminis* | 2 | (2) | 3 | (3) | 3 | (3) | 1 | (1) |  |  |  |  | 9 | (9) |
| *Agrilus hastulifer* | 7 | (6) | 3 | (3) | 2 | (2) |  |  |  |  |  |  | 12 | (11) |
| *Agrilus hyperici* |  |  |  |  | 14 | (14) | 13 | (11) | 8 | (8) | 7 | (7) | 42 | (40) |
| *Agrilus laticornis* | 29 | (25) | 21 | (19) | 31 | (26) | 41 | (35) | 1 | (1) |  |  | 123 | (106) |
| *Agrilus obscuricollis* * | 2 | (2) | 1 | (1) | 2 | (2) |  |  |  |  |  |  | 5 | (5) |
| *Agrilus olivicolor* | 8 | (6) | 8 | (6) | 5 | (5) | 1 | (1) |  |  |  |  | 22 | (18) |
| *Agrilus sulcicollis* * |  |  | 1 | (1) | 1 | (1) | 2 | (2) |  |  |  |  | 4 | (4) |
| *Agrilus viridis* * | 3 | (3) | 2 | (1) |  |  |  |  |  |  |  |  | 5 | (4) |
| *Coraebus undatus* | 4 | (4) |  |  | 2 | (2) | 1 | (1) |  |  |  |  | 7 | (7) |
| *Meliboeus fulgidicollis* | 6 | (5) |  |  | 1 | (0) | 2 | (2) |  |  |  |  | 9 | (7) |

The overall lifespan after capture was low and highly variable (Figure S3). The median was only 4 days, with a quarter of beetles living more than 6 days, 5 % (15 beetles) more than 13 days and 1 % (4 beetles) between 38 and 56 days.

Preflight mass differed significantly among species (Kruskal-Wallis, χ²(7) = 67.4, *p* < 0.001, n = 208). According to Dunn’s test, *C. undatus*, *A. hastulifer* and *A. graminis* were significantly heavier than the other species tested (Figure 2). *Coraebus undatus* was clearly out of category in terms of size and mass.

**Figure 2.**
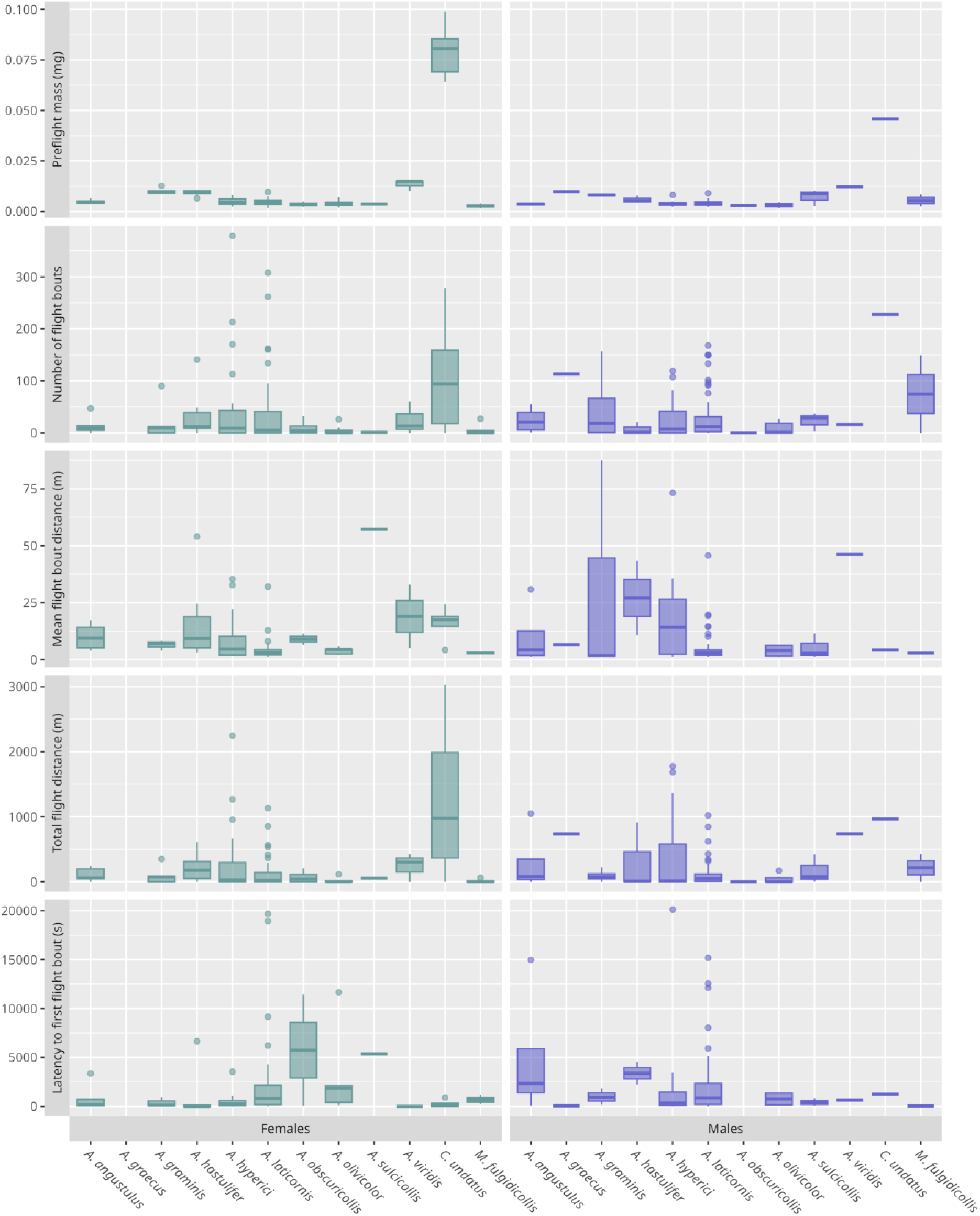
Values for a selection of response variables measured during the first (eight-hour) trial, per sex and species. Females: left (green), males: right (purple).

### General flight characteristics

First analyses were performed on first (eight-hour) trial data. All species and sexes combined, 222 individuals (123 females, 99 males) have been analysed, taking into account the removal of individuals for which some issue occurred during the trial. Main results are displayed in figure 2.

Regardless of the species, flight distance flown during the first trial was highly variable: 70 insects (31 %) did not fly, 44 % (98 beetles) flew between 1 m and 211 m, 25 % (56 beetles) flew more than 211 m, 10 % (23 beetles) more than 596 m, 5 % (12 beetles) more than 948 m, and 1 % (3 beetles) more than 2 138 m (Figure 3).The longest total distance flown within an eight-hour trial was 3 028 m for a female *C. undatus*, 2 246 m for a female *A. hyperici*, and 429 m for a male *M. fulgidicollis.* Remarkably, some individuals of *C. undatus* (two females) and *A. hyperici* (one female, four males) flew distances long enough to stand out from the right tail of the distributions, even after their first trial, and may be considered extreme (Figure 3). The flight speed of *Coraebus* and other species showed lower variation and was respectively 0.40 m/s ± 0.03 m/s and 0.26 m/s ± 0.13 m/s.

**Figure 3.**
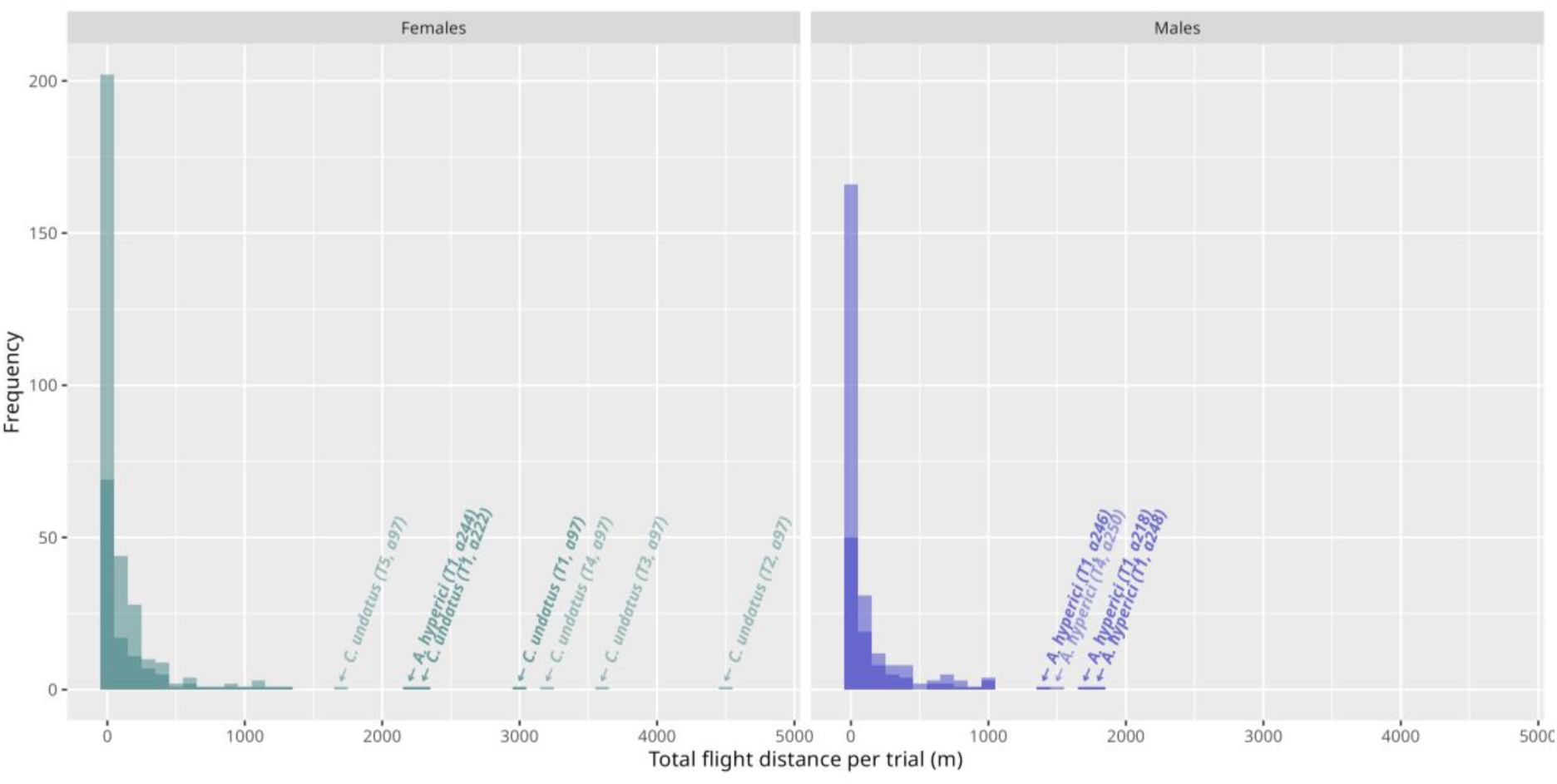
Distribution of total distance flown per trial depending on sex, all 12 species pooled together (n_females_ = 123, n_males_ = 99). Darker bars represent the first trial (eight hours) while lighter bars represent all consecutive trials (four hours), cumulated on top of the first trial. Individual flights ≥ 1300 m are marked with an arrow indicating the corresponding species, with the trial number and individual identifier in brackets.

Flight distance flown by the tested insects over their lifespan (i.e., including all the trials) varied also greatly. The median flight distance (Table 2) for *Agrilus* and *Meliboeus* was 53 m (115 m excluding the 28 % of non-flying insects); a quarter of the beetles flew more than 243 m (401 m), 10 % more than 730 m (1 039 m), and 1 % (3 beetles) flew between 2 890 m and 4 636 m (2 916 m and 4 636 m). Distances were longer for *C. undatus*: median distance of 967 m with one exceptional beetle above 16 km. Flown distance was globally linked with the number of trials (LM, p < 0.001, R² = 0.42, *C. undatus* excluded); no significant variation with species or sex had been observed.

**Table 2.** Median, mean and maximum total distance flown by the insects during their lifespan. First trial and Sum correspond to the data of the first eight-hour trial and the sum of all trials respectively.

| Species | Median total flight distance (m) |  |  |  | Mean total flight distance (m) |  |  |  | Highest total flight distance (m) |  |
| --- | --- | --- | --- | --- | --- | --- | --- | --- | --- | --- |
|  | With non-flying individuals |  | Without non-flying individuals |  | With non-flying individuals |  | Without non-flying individuals |  | First trial | Sum |
|  | First trial | Sum | First trial | Sum | First trial | Sum | First trial | Sum |  |  |
| *All with Coraebus undatus* | 40 | 54 | 95 | 119 | 200 ± 28 | 356 ± 74 | 290 ± 38 | 487 ± 100 | 3 028 | 16 097 |
| *All without Coraebus undatus* | 35 | 53 | 87 | 115 | 167 ± 22 | 274 ± 38 | 245 ± 31 | 379 ± 51 | 2 246 | 4 636 |
| *Agrilus angustulus* | 70 | 71 | 71 | 92 | 186 ± 100 | 205 ± 92 | 207 ± 109 | 225 ± 99 | 1 048 | 1 048 |
| *Agrilus graecus* | 738 | 738 | 738 | 738 | 738 | 738 | 738 | 738 | 738 | 738 |
| *Agrilus graminis* | 74 | 149 | 87 | 342 | 98 ± 39 | 663 ± 500 | 147 ± 47 | 995 ± 730 | 350 | 4 636 |
| *Agrilus hastulifer* | 143 | 182 | 212 | 212 | 245 ± 90 | 486 ± 189 | 300 ± 101 | 531 ± 202 | 910 | 2 135 |
| *Agrilus hyperici* | 25 | 56 | 316 | 460 | 341 ± 91 | 671 ± 152 | 546 ± 130 | 916 ± 189 | 2 246 | 2 989 |
| *Agrilus laticornis* | 35 | 53 | 67 | 95 | 121 ± 21 | 144 ± 21 | 169 ± 27 | 195 ± 26 | 1 132 | 1 132 |
| *Agrilus obscuricollis* | 0 | 0 | 144 | 144 | 58 ± 41 | 58 ± 41 | 144 ± 64 | 144 ± 64 | 208 | 208 |
| *Agrilus olivicolor* | 1 | 1 | 27 | 35 | 26 ± 11 | 28 ± 10 | 53 ± 19 | 52 ± 15 | 170 | 170 |
| *Agrilus sulcicollis* | 68 | 68 | 68 | 68 | 141 ± 96 | 141 ± 96 | 141 ± 96 | 141 ± 96 | 426 | 426 |
| *Agrilus viridis* | 364 | 782 | 428 | 894 | 367 ± 153 | 771 ± 277 | 489 ± 130 | 964 ± 256 | 739 | 1 644 |
| *Coraebus undatus* | 966 | 966 | 1 070 | 966 | 1 205 ± 410 | 3172 ± 2 168 | 1 405 ± 424 | 3172 ± 2 168 | 3 028 | 16 097 |
| *Meliboeus fulgidicollis* | 0 | 0 | 61 | 61 | 72 ± 60 | 79 ± 69 | 168 ± 131 | 211 ± 174 | 429 | 559 |

Regardless of their species, most tested individuals performed several short flight bouts within a single eight-hour trial (Figure 2). Flight bout number was highly variable: 25 or less for half of trials with flight activity, but more than 58 for a quarter (39 trials), and 16 trials (10 %) overreaching this number with more than 150 flight bouts over eight hours. Three trials even exceeded 300 flight bouts: two by females *A. hyperici* (407 and 379 flight bouts) and one by a female *A. laticornis* (308 flight bouts). The distance flown during a flight bout was also highly variable (median 2 m, 10% above 9.5 m, up to 653 m).

### Interspecific variation in flight performance

Among the four flight performance variables measured during the first (eight-hour) trial, two of them varied significantly among species (sex pooled): total flight distance (Kruskal-Wallis, χ²(7) = 20.4, *p* = 0.005, n = 208) and mean distance flown per flight bout (Kruskal-Wallis, χ²(7) = 23, *p* = 0.002, n = 208). However, Dunn’s tests highlighted only a few differences among groups: between *A. olivicolor* and *C. undatus* for the total flight distance, between *A. hastulifer* and *A. laticornis* for the mean distance flown per flight bouts. Latency to first flight bout (χ²(7) = 8.46, *p* = 0.29, n = 208) and number of flight bouts (χ²(7) = 13.7, *p* = 0.06, n = 208) were not influenced by species.

### Factors of intraspecific variation in flight performance

Intraspecific differences in the three most abundant species (*A. hyperici, A. laticornis* and *A. olivicolor*) demonstrated that endogenous factors influenced some of their flight capacities. Preflight mass positively affected total flight distance, as well as the number of flight bouts in *A. hyperici* and *A. laticornis*, but not in *A. olivicolor* (Table 3). Response variables showed no sex bias, as either fixed effects had no significant influence, or null models were better. However, marginal R² were weak and other factors likely have impacted flight performance. Only *A. hyperici* showed standing out marginal R² with number of moves (R² = 0.25) and total flight length (R² = 0.28) while every other R² did not exceed 0.13.

**Table 3.** LMER tests results for the effects of sex and preflight mass on a selection of response variables during the first (eight-hour) trial, with year of experiment as random effect. Response variables in each Linear Mixed-Effects Models were log-transformed. The reference level for the categorial factor ‘sex’ is female. ΔAICc = AICc (full model) – AICc (null model). The full model was considered to better explain the response variable than the null model when ΔAICc was lower than −2, and the opposite when ΔAICc was higher than −2. Conditional R² corresponds to the variance of both fixed and random effects while marginal R² considers only the variance of the fixed effect.

| Species | Variables | $\Delta AICc$ | Sex (female) | | | Mass | | | Cond. $R^2$ | Marg. $R^2$ |
| --- | --- | --- | --- | --- | --- | --- | --- | --- | --- | --- |
|  |  |  | Estimate | SE | t value | Estimate | SE | t value |  |  |
| *A. hyperici* | Total flight distance | -22.51 | 1.09 | 0.85 | 1.28 | 1 025.36 *** | 264.91 | 3.87 |  | 0.28 |
|  | Number of flight bouts | -19.98 | 0.45 | 0.61 | 0.73 | 679.24 *** | 189.78 | 3.58 |  | 0.25 |
|  | Mean flight bout distance | 8.62 | - | - | - | - | - | - | - | - |
|  | Latency to first flight bout | -11.33 | 0.85 | 1.07 | 0.79 | -308.19 | 369.73 | -0.83 | 0.09 | 0.07 |
| *A. laticornis* | Total flight distance | -17.81 | 0.81 | 0.44 | 1.85 | 461.98 ** | 150.70 | 3.07 | 0.14 | 0.09 |
|  | Number of flight bouts | -15.85 | 0.54 | 0.34 | 1.56 | 351.75 ** | 116.63 | 3.02 | 0.11 | 0.09 |
|  | Mean flight bout distance | -3.27 | 0.05 | 0.15 | 0.35 | 5.67 | 51.74 | 0.11 | 0.12 | 0.001 |
|  | Latency to first flight bout | -8.95 | -0.30 | 0.61 | -0.48 | 13.63 | 203.92 | 0.07 |  | 0.004 |
| *A. olivicolor* | Total flight distance | -10.82 | 1.58 | 0.97 | 1.63 | 456.42 | 346.36 | 1.31 | 0.45 | 0.12 |
|  | Number of flight bouts | -9.36 | 1.13 | 0.67 | 1.71 | 294.03 | 238.59 | 1.23 | 0.42 | 0.13 |
|  | Mean flight bout distance | 7.55 | - | - | - | - | - | - | - | - |
|  | Latency to first flight bout | 1.67 | - | - | - | - | - | - | - | - |

### Consecutive flight performance and patterns

In four-hour trials, the distribution of flight performance was similar to that of the first (eight-hour) one (Figure 3), but flight performance varied over consecutive trials. Data on *C. undatus* were excluded from these analyses, due to their specificities discussed below. Overall, raw flight performance decreased over consecutive trials (Figure 4 left, circle symbols; Kruskal-Wallis, χ²(9) = 46.2, p < 0.001, n = 501). Differences were mainly observed between the first trial and others. Extreme long distance flights (> 1 250 m, arbitrary threshold) only took place during the four first trials (and mainly in the first one). Flight performance increased in the late trials, when insects benefited from longer rest periods, but remained before first trials one (Figure 4 left, plus symbols). However, the global distribution showed that beetles flew a greater distance on their first trial, with no variation between subsequent trials (Figure 4 right; Kruskal-Wallis, χ²(9) = 140, p < 0.001, n = 501).

**Figure 4.**
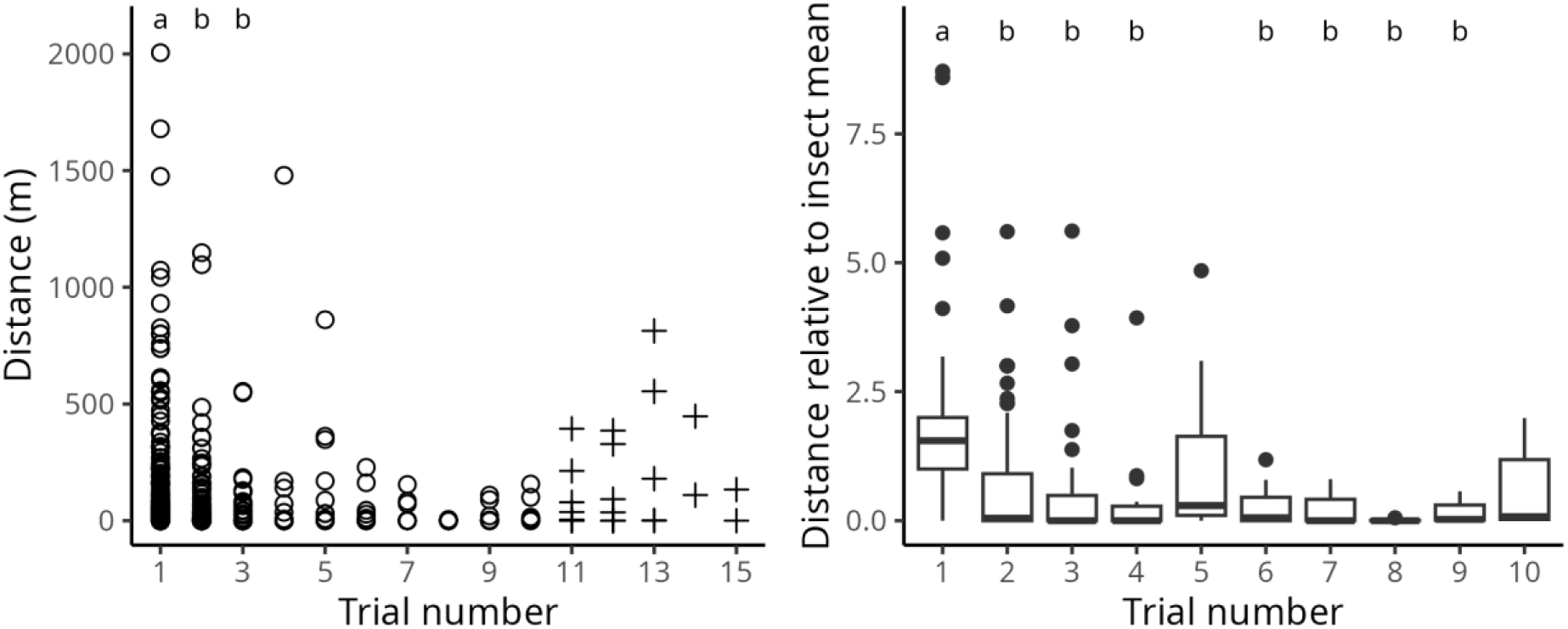
Changes in flight performance during consecutive trials. **Left:** Distribution of distance flown during four-hour trials across consecutive trials. **Right:** Distribution of relative flight performance across consecutive trials; for each beetle and trial, relative flight performance was computed as the flight distance during the trial divided by the mean flight distance during all trials. For both plots, data of the eight-hour trial was truncated to the first four hours. Unusual trial frequency data and data from *C. undatus* were discarded from analyses; however, late trial data are displayed on the left panel (plus symbols). Different letters above the data indicate a significant difference according to Dunn’s test.

Variable flight patterns over time were observed among species, with *A. hyperici, A. graminis,* and *C. undatus* standing out from the others because a fraction of their individuals remained alive and covered long distances after the first or even subsequent trials (Figures 5 and S4). No such pattern was observed in other species, either due to a generally low flight activity, or comparatively lower lifespan in rearing conditions, reducing the number of consecutive trials per individual. Regardless of species, 35 % of individuals were dead after the first trial, 70 % after the second and 86 % after the third. Most individuals (93 %) were dead after four trials. The vast majority of individuals that survived across multiple trials were unable to sustain long flight distances over time (mainly feeding the “[0]” category in figures 5 and S4), such as *A. laticornis* and *A. hastulifer*. However, some individuals diverged from this trend. A female *C. undatus* and a male *A. hyperici* flew long distances even after their first flight mill trial, as well as a female *A. hyperici* flying more than 800 m during its fifth trial (although not indicated with an arrow in figure 3; Figures 5 and S4). Remarkably, the extreme *C. undatus* female flew several kilometres in five consecutive trials, cumulating over 16 km of active flight after its capture.

**Figure 5.**
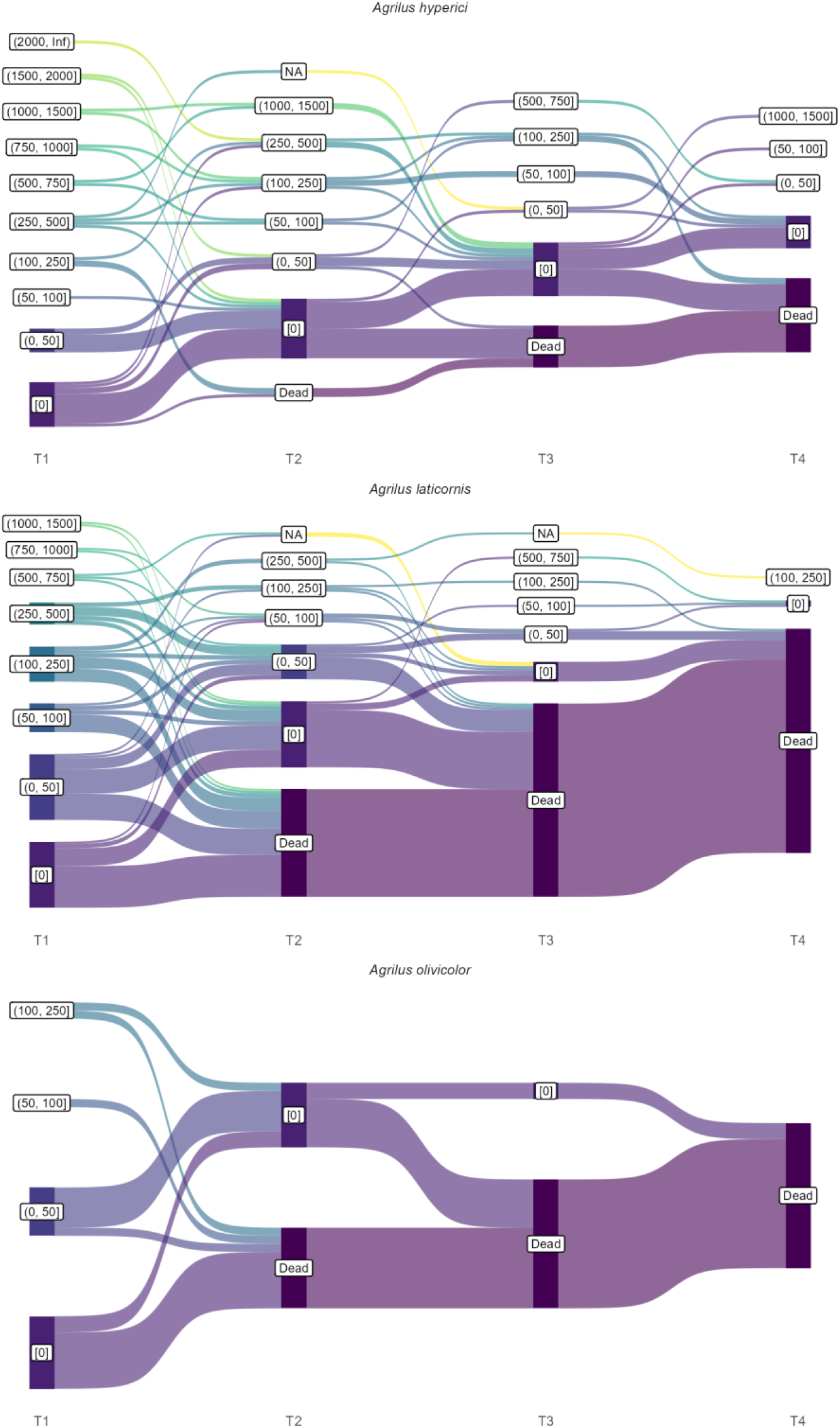
Individual flight trajectories across four consecutive trials in the three most numerous species (*A. laticornis*, *A. hyperici* and *A. olivicolor*), with the first trial spanning eight hours and the following four hours. Total flight distance is discretized in categories of increasing range to simplify individual trajectories. The NA category corresponds to missing data due to a technical issue during the trial (typically, a temporary escape of an individual).

## Discussion

### Flight performance and behaviour

This study allowed a first analysis of flight characteristics of several European Agrilinae, including flight performance and behaviour. It revealed several common patterns and some specificities among species. A clear observation was the high variation in most flight performance parameters among individuals. Prior history (including age and flight history) is a likely factor for this variation, but is unknown, albeit probably only part of the picture, since flight activity is typically known as an intrinsically variable trait among individuals and high inter-individual variability is common in flight mill experiments (Taylor et al., 2010; Sauvard et al., 2018). Flight performance decreased over the course of the trials, but changed little over a few ones; remarkably, the best flyer in *C. undatus* was able to fly long distances during several consecutive trials.

At the species level, the assessment of flight performance of several Agrilinae revealed rather homogeneous flight capacities among species, under the experimental conditions of tethered flights. Only the few *C. undatus* individuals presented substantial differences. Even if their number was too low to exhibit many significant results, these beetles actually performed some flights which largely exceeded the dispersal distance of any other species. However, this species is also much bigger than others. The huge intraspecific variation and the relatively low sample size, moreover unbalanced between species, had probably impeded identifying specific flight characteristics. *Coraebus undatus* excepted, the low variation in beetle size among the sampled species and the low flight activity of most tested individuals might have also reduced our ability to discriminate the results among species. Data available on flight capacities of other *Agrilus* species in similar experiment indicates better performance: median of 750 m for *A. planipennis* (Taylor et al., 2010) and mean between 200 to 1 800 m for *Agrilus auroguttatus* (Schaeffer), depending on conditions (Lopez et al., 2014). These values are rather similar to *C. undatus* performance, but, as this latter species, *A. planipennis* and *A. auroguttatus* are also bigger than the *Agrilus* and *Meliboeus* species we tested. From a more general perspective, Agrilinae performances are lower than those of some bigger beetles such as *Monochamus galloprovincialis* (Olivier) (2 km per day; David et al., 2014) and *Rhynchophorus ferrugineus* (Olivier) (2.6 km per day; Ávalos et al., 2014), and far from those of some long-flight insects such as hornets (median about 20 km per day; Sauvard et al., 2018).

At the trial level, most flying individuals showed no sustained flight, and instead completed multiple short flight bouts. This flying pattern could correspond to “routine flight” (Van Dyck & Baguette, 2005), typically associated with foraging or the search for a sexual partner or shelter. In contrast, these authors described longer flights as “dispersal flights”, presumably allowing insects to escape from bad conditions or predators, and were almost never observed during our experiment. However, opposing the two flight types may not always be relevant. In fact, our study showed that *Agrilus* species could spread by performing a series of short flights ultimately resulting in relatively long total distances covered, instead of single sustained flights (e.g., Ávalos et al., 2014).

Globally, tested beetles tended to be poor dispersers in the experimental setting, with most of them not taking off or flying over very short ranges (generally less than 200 m). Only *C. undatus* seemed to fly significantly longer distances (median of 1 000 m). This observation may suggest resident strategies, i.e., individuals that tend to stay in the range of their emergence area. One third of beetles did not fly at all. This is congruent with studies on other beetle species, which found a similar percentage of non-flyers, such as for *Ips sexdentatus* Boern (Jactel, 1993) and *R. ferrugineus* (Ávalos et al., 2014). These authors hypothesised that such a high proportion of non-flying insects might be due to the tethered of the beetles and to the flight mill itself, or be a consequence of the individual incapacity to fly. As some non-flying beetles in a trial also demonstrated being able to fly during a later trial, not flying was probably due to a combination of behavioural variations, propensity to take-off, and constraints of the flight mills, but not solely the latter. Nevertheless, as flight performance distribution had a very long tail, some insects largely exceeded the median performance of their conspecifics and could be considered good dispersers. As such, they may perform better at reaching new distant habitats if enough patches of viable hosts are available on their path (Lopez et al., 2014), which is normally the case for oak-associated species in European temperate areas. Within *Agrilus* species, an *A. hyperici* appeared to be the most active disperser; interestingly, this species was the only one that was not associated with forest habitats in our sample, and their relatively high performance could be related to its more scattered herbaceous habitat. *A. hyperici* was introduced in western Canada during the last century and, even though no accurate data is available, it seems to have largely spread in this region since; however, this took several decades despite active human assistance (Gov. of British Columbia, 2018). This observation *in natura* is consistent with modest flight capacities in this species. Alongside their flight capacities, it is possible that small Agrilinae species occasionally take advantage of winds to maximize dispersal, as observed in some bark beetles up to several tens of kilometres (Nilssen, 1984). Moreover, the actual realized dispersal in Agrilinae could result from the combination of short-distance flight and human-assisted transportation (i.e., with wood material), sometimes over long distances, as observed in *A. planipennis* (Muirhead et al., 2006), *Agrilus bilineatus* (Weber) (Baranchikov et al., 2019b) and related species (e.g., Wu et al., 2017; Ruzzier et al., 2023).

### No sexual dimorphism in flight performance

We expected greater flight performance in females than males, as already demonstrated with mated females of *A. planipennis* (Taylor et al., 2010). Yet, sex did not impact any flight variables in the overall sample. Those results are concordant, however, with observations on *A. auroguttatus* whose flight capacities were not affected by sex either (Lopez et al., 2014). Lack of sexual dimorphism in flight performance seems then frequent in Agrilinae species. Both sexes are suspected to participate in dispersal in the tested species. Males are known to use flight to actively search for females, while mated females typically disperse to find a suitable host for egg laying (Chamorro et al., 2015), and both behaviours may contribute to species dispersal. However, it is rather considered that females are the ones which are able to disperse far away from their host when necessary while males stay close to their natal or maturation area (Rodriguez-Soana et al., 2007; Mercader et al., 2009; Taylor et al., 2010).

### Importance of body mass on flight performance

Body mass is known to be positively correlated with flight performance in many insect species, and is generally considered to reflect the available fuel load (Minter et al., 2018). Such a relation was observed in the majority of the species we tested, including the best flyer (*C. undatus*) and the two most numerous species (*A. laticornis* and *A. hyperici*). Relation to body mass seemed to differ between these groups; in *C. undatus*, heavier individuals tend to fly immediately and to perform longer bouts; in the two *Agrilus*, heavier individuals tend, on the contrary, to increase the number of bouts. In contrast to these species, performance of other ones, including the third most numerous one (*A. olivicolor*), was not impacted by preflight mass, which corroborates what was observed in some other *Agrilus* species, i.e., *A. auroguttatus* (Lopez et al., 2014) and *A. planipennis* (Taylor et al., 2010). This difference could be related to the relative importance of reserves *vs.* dead weight (mainly structure) in mass variation, but also to insect strategy (noticeably, they could mainly fly when reserves are high, or, on the contrary, when they are low, to forage in order to increase them). It should be noted, however, that even when preflight mass was related to flight performance, it explains only a small part of its variation. Other factors should be considered, and the effect of some of them, diet, mated status and age, has been demonstrated in other *Agrilus* species (Taylor et al., 2010; Lopez et al., 2014). Future studies should include these factors to further investigate Agrilinae flight performance. Nevertheless, our results on species of different body mass (*Agrilus* and *Meliboeus* species *vs. C. undatus*) and previous literature on *A. planipennis* and *A. auroguttatus* suggest that the positive correlation between body mass and flight performance stands among Agrilinae species.

### Limitations of the study and interpretation caveats

The sampling design based on interception traps allowed capturing multiple species associated with oak forests, and not only oak borers. However, it prevented controlling for additional factors pertaining to the prior experience of individuals (flight, diet, mated status) and inherently biased captures towards dispersing individuals. This may cause either (1) undersampling of the most resident individuals, even more so than those identified in this study, or (2) sampling of individuals that already spent energy on dispersal before the experiment, or a combination of both. Together with the limited number of captured individuals for some species and the impossibility to control some factors, this warrants using a different sampling strategy in further studies focusing on only oak-borers. We used dry collectors in green multi-funnel traps which are a relevant method for Buprestidae (Allison & Redak, 2017). However, our sampling design likely let a fraction of individuals escape even if the collector was emptied every workday, possibly those showing the highest locomotor activity. Indeed, similar traps but equipped with wet collectors were installed for another study nearby one of the sampling sites, and captured a more abundant and richer community of beetles (A. Sallé, pers. comm.). Emergence traps can yield very good results (e.g., Taylor et al., 2010; Lopez et al., 2014 for borers), but would have restricted the sample to borers of the equipped hosts only, instead of targeting all circulating Buprestidae in oak forests.

The intrinsic dispersal potential of a flying insect species is generally determined mainly by the overall flight capacity of its individuals throughout their life, more specifically that of its best flyers. In the Agrilinae species studied here (except perhaps *C. undatus*), this capacity appeared low, with about 25% of insects not flying in the experimental conditions they were subjected to, and actively flying individuals rarely overreaching one kilometre. As flight performance appeared to be rather maintained over consecutive trials, flight capacity was highly dependent on lifespan. However, the beetles proved to be difficult to maintain alive in rearing conditions, with a median lifespan of four days across all species. As a comparison, *A. planipennis* was reported to live between 2 and 17 weeks under laboratory conditions, with an average of 7 weeks for males and 9 weeks for females (Chamorro et al., 2015 and references within), albeit figures indicated a considerable variability in lifespan in this species as well. Given our observations of limited leaf damage and faeces accumulation in the rearing vials, we suspect insects failed to feed properly in the conditions they were offered, for unknown reasons. Our method was nevertheless the same one used successfully by other studies of flight performance of *Agrilus* spp. (Taylor et al., 2010; Lopez et al., 2014). Combined effects of starvation and daily flight trials in flight mills likely impacted survival severely in the present experiment, ultimately reducing lifespan and underestimating the overall flight capacity of these species. Moreover, lifespan values are to be taken with caution since the age of individuals at the time of their capture remains unknown.

Flight mills under controlled conditions are a proven method to assess the flight potential of insects (e.g., Sauvard et al., 2018 and references therein). Since the method can easily be standardized under controlled conditions, flight mills are especially valuable to compare species or investigate the influence of factors on flight activity. However, tethered flight performance cannot be considered a direct measurement of free flight or natural flight (Ribak et al., 2017; Sauvard et al., 2018; Naranjo, 2019), and cautious interpretation is advised. Flight mills create artificial conditions and could both hinder take-off (inertia) or promote flight (support making the insects weightless, inertia of the rotating arm). The absence of a moving natural landscape in the vision of the specimens is likely to influence their locomotor behaviour, and the influence of sensed angular velocity or weightlessness on tethered insects’ propensity to fly is unknown. Tethered flight experiments have already been suspected of under- or overestimating flight abilities due to the lack of tarsal contact and the absence of a perch for taking off or landing (Minter et al., 2018) or because of the device itself (Taylor et al., 2010). Yet, they remain a highly relevant and standard method to compare conspecifics or sympatric species in a given set of conditions to approach their motivation to disperse under those settings, or even physiological expenditures in flight dispersal when their flight propensity is high. In other respects, flight mills likely limited the flight speed in our experiment, as was found in another flight mill study on *A. planipennis* with a three-fold difference between free and tethered flight speeds (Taylor et al., 2010).

## Conclusions and perspectives

This work provided a first comparative approach on the flight behaviour and performance in several Agrilinae species associated with oak forests. It showed several common traits among the focal species, namely (1) the considerable inter-individual variation, (2) the low average flight performance compared to other insect species using similar experimental designs, and (3) the relative homogeneity of this pattern among most of the species investigated, despite the probable existence of different flight behaviour among species as seen with long sustained flights in *C. undatus*. This provides insights into the poorly understood dispersal ecology of oak-associated Agrilinae, suggesting a generally low average dispersal propensity and the importance of scarce events carried by a few extreme individuals in shaping the ultimate colonization and spread patterns at the population and species levels.

While the present work aimed at investigating flight in Agrilinae using a multispecific approach that, to our knowledge, has rarely been attempted with flight mill experiments, an oak-borer species considered a key aggravating agent of oak decline in Europe is missing in our sampling: *A. biguttatus*. This beetle is currently expanding its range with climate change (Pederson & Jørum, 2009; Brown et al., 2015) and genetic molecular markers have highlighted population differentiation at the European scale (Le Souchu et al. submitted). Yet, little is known about the dispersal ability of this species, let alone its variation among populations. Flight mills may prove a relevant method to better understand the genetic structure and movements of populations in this species, especially as it is considered a major threat outside its current range (e.g., in North America, USDA-APHIS, 2011). As the biggest European *Agrilus* species, at a size similar to that of *A. planipennis* (Volkovitsh et al., 2019) in which flight ability is described in most details among *Agrilus* species, comparing both species would be insightful to better understand the spread potential of *A. biguttatus*.

## Appendices

The supplementary material is available in the open repository Data Recherche Gouv at https://doi.org/10.57745/TY9KLG.

## Acknowledgements

We are thankful for the technical assistance offered by Maxime Stanislawek, Clément Beaumont and Mathieu Plateau. A preprint version of this article has been peer-reviewed and recommended by PCI Zool (https://doi.org/10.24072/pci.zool.100354).

## Data, scripts, code, and supplementary information availability

The data and code that support the findings of this study are openly available in three repositories. The main one, at https://doi.org/10.57745/TY9KLG, includes the material specific to the present study: supplementary data, videos of insects flying in the rotational flight mills, raw log files from the flight mills, associated processed data and final *R* scripts. Git-versioned *R* scripts are also available at https://forgemia.inra.fr/mathieu.laparie/agrilusflight. The source of the *Crystal* program used to monitor flight mills and produce log files is available at https://doi.org/10.57745/DIWSWF. The *Ruby* pipelines used to process log files into flight data are available at https://doi.org/10.57745/YLMCNU.

