## Supplementary material for "Intra- and interspecific variations in flight performance of oak-associated Agrilinae (Coleoptera: Buprestidae) using computerised flight mills": S1

Appendices

**Table S1.** List of Agrilinae individuals tested in 2021, 2022 and 2023 using computerised flight mills. Lat. and Lon. are latitude and longitude respectively, and are displayed in WGS 84, DDD, while M and F correspond to male and female.

| **Insect ID** | **Species** | **Sex** | **Trapping site** | **Lat.** | **Lon.** | **Trapping date** | **Year** | **Death date** | **has been lost** |
| --- | --- | --- | --- | --- | --- | --- | --- | --- | --- |
| **a1** | *A. laticornis* | M | University of Orléans | 47.85 | 1.94 | 06-23-2021 | 2021 | 06-26-2021 |  |
| **a2** | *A. laticornis* | M | University of Orléans | 47.85 | 1.94 | 06-23-2021 | 2021 | 06-26-2021 |  |
| **a3** | *A. laticornis* | F | University of Orléans | 47.85 | 1.94 | 06-24-2021 | 2021 | 06-26-2021 |  |
| **a4** | *M. fulgidicollis* | F | University of Orléans | 47.85 | 1.94 | 06-24-2021 | 2021 | 06-26-2021 |  |
| **a5** | *A. hastulifer* | M | University of Orléans | 47.85 | 1.94 | 06-18-2021 | 2021 | 06-29-2021 |  |
| **a6** | *A. olivicolor* | F | University of Orléans | 47.85 | 1.94 | 06-18-2021 | 2021 | 06-29-2021 |  |
| **a7** |  |  | University of Orléans | 47.85 | 1.94 | 06-22-2021 | 2021 | 06-28-2021 | yes |
| **a8** | *A. laticornis* | F | University of Orléans | 47.85 | 1.94 | 06-28-2021 | 2021 | 06-30-2021 |  |
| **a9** | *A. hastulifer* | F | University of Orléans | 47.85 | 1.94 | 06-28-2021 | 2021 | 08-05-2021 |  |
| **a10** | *A. viridis* | M | University of Orléans | 47.85 | 1.94 | 06-28-2021 | 2021 | 06-30-2021 |  |
| **a11** | *A. olivicolor* | M | University of Orléans | 47.85 | 1.94 | 06-28-2021 | 2021 | 06-30-2021 |  |
| **a12** |  |  | INRAE | 47.83 | 1.91 | 06-29-2021 | 2021 | 06-30-2021 | yes |
| **a13** |  |  | University of Orléans | 47.85 | 1.94 | 06-29-2021 | 2021 | 06-30-2021 | yes |
| **a14** | *A. laticornis* | F | University of Orléans | 47.85 | 1.94 | 06-29-2021 | 2021 | 07-01-2021 |  |
| **a15** | *A. laticornis* | M | University of Orléans | 47.85 | 1.94 | 06-29-2021 | 2021 | 07-01-2021 |  |
| **a16** | *A. olivicolor* | F | University of Orléans | 47.85 | 1.94 | 06-30-2021 | 2021 | 07-02-2021 |  |
| **a17** | *A. laticornis* | M | INRAE | 47.83 | 1.91 | 07-01-2021 | 2021 | 07-03-2021 |  |
| **a18** | *A. laticornis* | M | University of Orléans | 47.85 | 1.94 | 07-01-2021 | 2021 | 07-02-2021 |  |
| **a19** | *A. graminis* | M | University of Orléans | 47.85 | 1.94 | 07-01-2021 | 2021 | 08-06-2021 |  |
| **a20** | *A. olivicolor* | M | University of Orléans | 47.85 | 1.94 | 07-01-2021 | 2021 | 07-02-2021 |  |
| **a21** | *A. obscuricollis* | M | INRAE | 47.83 | 1.91 | 07-01-2021 | 2021 | 07-02-2021 |  |
| **a22** | *A. laticornis* | F | INRAE | 47.83 | 1.91 | 07-02-2021 | 2021 | 07-06-2021 |  |
| **a23** | *A. laticornis* | M | INRAE | 47.83 | 1.91 | 07-02-2021 | 2021 | 07-05-2021 |  |
| **a24** |  |  | INRAE | 47.83 | 1.91 | 07-05-2021 | 2021 | 07-06-2021 | yes |
| **a25** | *A. laticornis* | F | INRAE | 47.83 | 1.91 | 07-05-2021 | 2021 | 07-07-2021 |  |
| **a26** | *A. olivicolor* | F | University of Orléans | 47.85 | 1.94 | 07-05-2021 | 2021 | 07-09-2021 |  |
| **a27** | *A. hastulifer* | M | University of Orléans | 47.85 | 1.94 | 07-05-2021 | 2021 | 07-09-2021 |  |
| **a28** | *A. olivicolor* | M | University of Orléans | 47.85 | 1.94 | 07-05-2021 | 2021 | 07-08-2021 |  |
| **a29** | *A. graminis* | M | University of Orléans | 47.85 | 1.94 | 07-05-2021 | 2021 | 07-09-2021 |  |
| **a30** | *A. laticornis* | M | University of Orléans | 47.85 | 1.94 | 07-05-2021 | 2021 | 07-08-2021 |  |
| **a31** | *A. viridis* | F | INRAE | 47.83 | 1.91 | 07-06-2021 | 2021 | 07-12-2021 |  |
| **a32** | *A. laticornis* | M | University of Orléans | 47.85 | 1.94 | 07-06-2021 | 2021 | 07-12-2021 |  |
| **a33** | *A. laticornis* | F | University of Orléans | 47.85 | 1.94 | 07-06-2021 | 2021 | 07-08-2021 |  |
| **a34** |  |  | University of Orléans | 47.85 | 1.94 | 07-06-2021 | 2021 | 07-08-2021 | yes |
| **a35** | *A. laticornis* | F | University of Orléans | 47.85 | 1.94 | 07-06-2021 | 2021 | 07-09-2021 |  |
| **a36** | *A. laticornis* | M | University of Orléans | 47.85 | 1.94 | 07-07-2021 | 2021 | 07-09-2021 |  |
| **a37** | *A. laticornis* | M | University of Orléans | 47.85 | 1.94 | 07-07-2021 | 2021 | 07-12-2021 |  |
| **a38** | *A. laticornis* | F | University of Orléans | 47.85 | 1.94 | 07-07-2021 | 2021 | 07-08-2021 |  |
| **a39** | *A. olivicolor* | M | University of Orléans | 47.85 | 1.94 | 07-07-2021 | 2021 | 07-12-2021 |  |
| **a40** | *A. laticornis* | F | INRAE | 47.83 | 1.91 | 07-08-2021 | 2021 | 07-12-2021 |  |
| **a41** | *A. laticornis* | M | INRAE | 47.83 | 1.91 | 07-08-2021 | 2021 | 07-12-2021 |  |
| **a42** | *M. fulgidicollis* | F | University of Orléans | 47.85 | 1.94 | 07-08-2021 | 2021 | 07-12-2021 |  |
| **a43** | *A. laticornis* | M | University of Orléans | 47.85 | 1.94 | 07-08-2021 | 2021 | 07-12-2021 |  |
| **a44** | *A. laticornis* | F | University of Orléans | 47.85 | 1.94 | 07-08-2021 | 2021 | 07-12-2021 |  |
| **a45** | *A. olivicolor* | F | University of Orléans | 47.85 | 1.94 | 07-08-2021 | 2021 | 07-12-2021 |  |
| **a46** | *A. laticornis* | F | INRAE | 47.83 | 1.91 | 07-08-2021 | 2021 | 07-12-2021 |  |
| **a47** | *A. laticornis* | M | University of Orléans | 47.85 | 1.94 | 07-09-2021 | 2021 | 07-13-2021 |  |
| **a48** | *A. laticornis* | F | University of Orléans | 47.85 | 1.94 | 07-09-2021 | 2021 | 07-13-2021 |  |
| **a49** | *A. hastulifer* | F | University of Orléans | 47.85 | 1.94 | 07-12-2021 | 2021 | 07-19-2021 |  |
| **a50** | *A. sulcicollis* | M | University of Orléans | 47.85 | 1.94 | 07-12-2021 | 2021 | 07-15-2021 |  |
| **a51** | *A. laticornis* | M | University of Orléans | 47.85 | 1.94 | 07-12-2021 | 2021 | 07-15-2021 |  |
| **a52** | *M. fulgidicollis* | F | University of Orléans | 47.85 | 1.94 | 07-12-2021 | 2021 | 07-15-2021 |  |
| **a53** | *A. hastulifer* | F | University of Orléans | 47.85 | 1.94 | 07-12-2021 | 2021 | 07-19-2021 |  |
| **a54** | *A. laticornis* | M | University of Orléans | 47.85 | 1.94 | 07-12-2021 | 2021 | 07-16-2021 |  |
| **a55** | *M. fulgidicollis* | F | INRAE | 47.83 | 1.91 | 07-12-2021 | 2021 | 07-19-2021 |  |
| **a56** | *A. laticornis* | M | University of Orléans | 47.85 | 1.94 | 07-13-2021 | 2021 | 07-19-2021 |  |
| **a57** |  |  | University of Orléans | 47.85 | 1.94 | 07-13-2021 | 2021 | 07-15-2021 | yes |
| **a58** | *A. laticornis* | M | University of Orléans | 47.85 | 1.94 | 07-15-2021 | 2021 | 07-19-2021 |  |
| **a59** | *A. olivicolor* | M | University of Orléans | 47.85 | 1.94 | 07-15-2021 | 2021 | 07-19-2021 |  |
| **a60** | *A. olivicolor* | M | University of Orléans | 47.85 | 1.94 | 07-15-2021 | 2021 | 07-19-2021 |  |
| **a61** | *A. laticornis* | M | University of Orléans | 47.85 | 1.94 | 07-15-2021 | 2021 | 07-19-2021 |  |
| **a62** | *A. angustulus* | F | University of Orléans | 47.85 | 1.94 | 07-16-2021 | 2021 | 07-20-2021 |  |
| **a63** | *A. laticornis* | F | University of Orléans | 47.85 | 1.94 | 07-16-2021 | 2021 | 07-19-2021 |  |
| **a64** | *A. viridis* | F | INRAE | 47.83 | 1.91 | 07-16-2021 | 2021 | 07-19-2021 |  |
| **a65** | *A. graecus* | M | University of Orléans | 47.85 | 1.94 | 07-19-2021 | 2021 | 07-22-2021 |  |
| **a66** | *A. olivicolor* | F | University of Orléans | 47.85 | 1.94 | 07-19-2021 | 2021 | 07-22-2021 |  |
| **a67** | *A. olivicolor* | F | University of Orléans | 47.85 | 1.94 | 07-19-2021 | 2021 | 07-22-2021 |  |
| **a68** | *A. olivicolor* | F | University of Orléans | 47.85 | 1.94 | 07-19-2021 | 2021 | 07-22-2021 |  |
| **a69** | *A. laticornis* | F | University of Orléans | 47.85 | 1.94 | 07-19-2021 | 2021 | 07-22-2021 |  |
| **a70** | *M. fulgidicollis* | F | INRAE | 47.83 | 1.91 | 07-20-2021 | 2021 | 07-26-2021 |  |
| **a71** | *A. laticornis* | M | University of Orléans | 47.85 | 1.94 | 07-21-2021 | 2021 | 07-23-2021 |  |
| **a72** | *A. laticornis* | F | University of Orléans | 47.85 | 1.94 | 07-21-2021 | 2021 | 07-26-2021 |  |
| **a73** | *A. laticornis* | F | University of Orléans | 47.85 | 1.94 | 07-21-2021 | 2021 | 07-26-2021 |  |
| **a74** | *A. laticornis* | F | University of Orléans | 47.85 | 1.94 | 07-21-2021 | 2021 | 07-26-2021 |  |
| **a75** | *A. laticornis* | F | University of Orléans | 47.85 | 1.94 | 07-21-2021 | 2021 | 07-26-2021 |  |
| **a76** | *A. laticornis* | M | University of Orléans | 47.85 | 1.94 | 07-21-2021 | 2021 | 07-23-2021 |  |
| **a77** | *A. olivicolor* | M | University of Orléans | 47.85 | 1.94 | 07-21-2021 | 2021 | 07-23-2021 |  |
| **a78** | *A. hastulifer* | F | INRAE | 47.83 | 1.91 | 07-22-2021 | 2021 | 08-02-2021 |  |
| **a79** | *A. laticornis* | M | INRAE | 47.83 | 1.91 | 07-22-2021 | 2021 | 07-26-2021 |  |
| **a80** | *A. laticornis* | F | INRAE | 47.83 | 1.91 | 07-22-2021 | 2021 | 07-26-2021 |  |
| **a81** | *A. laticornis* | F | INRAE | 47.83 | 1.91 | 07-22-2021 | 2021 | 07-26-2021 |  |
| **a82** | *A. laticornis* | F | University of Orléans | 47.85 | 1.94 | 07-22-2021 | 2021 | 07-26-2021 |  |
| **a83** | *A. laticornis* | F | University of Orléans | 47.85 | 1.94 | 07-22-2021 | 2021 | 07-26-2021 |  |
| **a84** | *A. laticornis* | F | University of Orléans | 47.85 | 1.94 | 07-22-2021 | 2021 | 07-26-2021 |  |
| **a85** | *A. obscuricollis* | F | University of Orléans | 47.85 | 1.94 | 07-22-2021 | 2021 | 07-26-2021 |  |
| **a86** | *A. obscuricollis* | F | University of Orléans | 47.85 | 1.94 | 07-22-2021 | 2021 | 07-26-2021 |  |
| **a87** | *A. laticornis* | F | University of Orléans | 47.85 | 1.94 | 07-22-2021 | 2021 | 07-26-2021 |  |
| **a88** | *C. undatus* | F | University of Orléans | 47.85 | 1.94 | 07-22-2021 | 2021 | 07-26-2021 |  |
| **a89** | *A. laticornis* | F | University of Orléans | 47.85 | 1.94 | 07-23-2021 | 2021 | 07-26-2021 |  |
| **a90** | *A. hastulifer* | M | University of Orléans | 47.85 | 1.94 | 07-23-2021 | 2021 | 07-27-2021 |  |
| **a91** | *A. viridis* | M | University of Orléans | 47.85 | 1.94 | 07-27-2021 | 2021 | 07-30-2021 |  |
| **a92** | *A. laticornis* | F | University of Orléans | 47.85 | 1.94 | 07-27-2021 | 2021 | 07-29-2021 |  |
| **a93** | *C. undatus* | F | University of Orléans | 47.85 | 1.94 | 07-27-2021 | 2021 | 07-30-2021 |  |
| **a94** | *C. undatus* | F | University of Orléans | 47.85 | 1.94 | 07-27-2021 | 2021 | 07-29-2021 |  |
| **a95** | *A. laticornis* | F | University of Orléans | 47.85 | 1.94 | 07-29-2021 | 2021 | 08-02-2021 |  |
| **a96** | *M. fulgidicollis* | F | University of Orléans | 47.85 | 1.94 | 07-29-2021 | 2021 | 07-30-2021 |  |
| **a97** | *C. undatus* | F | University of Orléans | 47.85 | 1.94 | 07-29-2021 | 2021 | 08-04-2021 |  |
| **a98** | *A. hastulifer* | F | University of Orléans | 47.85 | 1.94 | 07-30-2021 | 2021 | 08-04-2021 |  |
| **a99** | *A. laticornis* | F | University of Orléans | 47.85 | 1.94 | 08-02-2021 | 2021 | 08-04-2021 |  |
| **a100** | *A. hastulifer* | F | INRAE | 47.83 | 1.91 | 08-02-2021 | 2021 | 08-05-2021 |  |
| **a101** | *A. graminis* | F | INRAE | 47.83 | 1.91 | 08-02-2021 | 2021 | 09-13-2021 |  |
| **a102** | *A. graminis* | F | INRAE | 47.83 | 1.91 | 08-03-2021 | 2021 | 08-16-2021 |  |
| **a103** | *A. graminis* | M | INRAE | 47.83 | 1.91 | 08-03-2021 | 2021 | 08-16-2021 |  |
| **a104** | *A. olivicolor* | M | University of Orléans | 47.85 | 1.94 | 08-03-2021 | 2021 | 08-05-2021 |  |
| **a105** | *A. viridis* | F | University of Orléans | 47.85 | 1.94 | 08-03-2021 | 2021 | 08-09-2021 |  |
| **a106** | *A. laticornis* | F | University of Orléans | 47.85 | 1.94 | 08-03-2021 | 2021 | 08-06-2021 |  |
| **a107** | *A. olivicolor* | F | University of Orléans | 47.85 | 1.94 | 08-03-2021 | 2021 | 08-05-2021 |  |
| **a108** | *A. hastulifer* | F | University of Orléans | 47.85 | 1.94 | 08-05-2021 | 2021 | 09-16-2021 |  |
| **a109** | *A. angustulus* | M | INRAE | 47.83 | 1.91 | 05-09-2022 | 2022 | 05-17-2022 |  |
| **a110** | *A. sulcicollis* | M | INRAE | 47.83 | 1.91 | 05-10-2022 | 2022 | 05-17-2022 |  |
| **a111** | *A. laticornis* | M | INRAE | 47.83 | 1.91 | 05-12-2022 | 2022 | 05-18-2022 |  |
| **a112** | *A. laticornis* | M | INRAE | 47.83 | 1.91 | 05-16-2022 | 2022 | 05-19-2022 |  |
| **a113** | *A. obscuricollis* | F | University of Orléans | 47.85 | 1.94 | 05-16-2022 | 2022 | 05-19-2022 |  |
| **a114** | *A. laticornis* | F | INRAE | 47.83 | 1.91 | 05-19-2022 | 2022 | 05-25-2022 |  |
| **a115** | *A. laticornis* | M | INRAE | 47.83 | 1.91 | 05-19-2022 | 2022 | 05-23-2022 |  |
| **a116** | *A. laticornis* | F | INRAE | 47.83 | 1.91 | 05-19-2022 | 2022 | 05-23-2022 |  |
| **a117** | *A. obscuricollis* | F | University of Orléans | 47.85 | 1.94 | 05-19-2022 | 2022 | 05-24-2022 |  |
| **a118** | *A. laticornis* | F | University of Orléans | 47.85 | 1.94 | 05-19-2022 | 2022 | 05-23-2022 |  |
| **a119** | *A. laticornis* | F | INRAE | 47.83 | 1.91 | 05-20-2022 | 2022 | 05-24-2022 |  |
| **a120** | *A. laticornis* | M | INRAE | 47.83 | 1.91 | 05-20-2022 | 2022 | 05-25-2022 |  |
| **a121** | *A. laticornis* | M | INRAE | 47.83 | 1.91 | 05-20-2022 | 2022 | 05-25-2022 |  |
| **a122** | *A. laticornis* | M | INRAE | 47.83 | 1.91 | 05-20-2022 | 2022 | 05-25-2022 |  |
| **a123** | *A. angustulus* | M | University of Orléans | 47.85 | 1.94 | 05-30-2022 | 2022 | 06-03-2022 |  |
| **a124** | *A. laticornis* | M | INRAE | 47.83 | 1.91 | 05-31-2022 | 2022 | 06-07-2022 |  |
| **a125** | *A. laticornis* | M | INRAE | 47.83 | 1.91 | 06-01-2022 | 2022 | 06-07-2022 |  |
| **a126** | *A. laticornis* | M | INRAE | 47.83 | 1.91 | 06-01-2022 | 2022 | 06-07-2022 |  |
| **a127** | *A. laticornis* | M | INRAE | 47.83 | 1.91 | 06-01-2022 | 2022 | 06-07-2022 |  |
| **a128** | *A. laticornis* | M | INRAE | 47.83 | 1.91 | 06-01-2022 | 2022 | 06-03-2022 |  |
| **a129** | *A. laticornis* | M | INRAE | 47.83 | 1.91 | 06-01-2022 | 2022 | 06-07-2022 |  |
| **a130** |  |  | INRAE | 47.83 | 1.91 | 06-01-2022 | 2022 | 06-02-2022 | yes |
| **a131** | *A. laticornis* | F | INRAE | 47.83 | 1.91 | 06-01-2022 | 2022 | 06-07-2022 |  |
| **a132** | *A. laticornis* | F | INRAE | 47.83 | 1.91 | 06-01-2022 | 2022 | 06-07-2022 |  |
| **a133** | *M. fulgidicollis* | M | University of Orléans | 47.85 | 1.94 | 06-01-2022 | 2022 | 06-07-2022 |  |
| **a134** | *A. sulcicollis* | F | University of Orléans | 47.85 | 1.94 | 06-01-2022 | 2022 | 06-03-2022 |  |
| **a135** | *A. sulcicollis* | M | University of Orléans | 47.85 | 1.94 | 06-01-2022 | 2022 | 06-07-2022 |  |
| **a136** | *A. olivicolor* | F | University of Orléans | 47.85 | 1.94 | 06-01-2022 | 2022 | 06-08-2022 |  |
| **a137** | *A. laticornis* | M | INRAE | 47.83 | 1.91 | 06-02-2022 | 2022 | 06-07-2022 |  |
| **a138** | *A. laticornis* | M | INRAE | 47.83 | 1.91 | 06-02-2022 | 2022 | 06-07-2022 |  |
| **a139** | *A. laticornis* | M | INRAE | 47.83 | 1.91 | 06-02-2022 | 2022 | 06-07-2022 |  |
| **a140** | *A. laticornis* | M | INRAE | 47.83 | 1.91 | 06-02-2022 | 2022 | 06-08-2022 |  |
| **a141** | *A. laticornis* | M | INRAE | 47.83 | 1.91 | 06-02-2022 | 2022 | 06-10-2022 |  |
| **a142** | *A. laticornis* | M | INRAE | 47.83 | 1.91 | 06-02-2022 | 2022 | 06-08-2022 |  |
| **a143** | *A. laticornis* | M | INRAE | 47.83 | 1.91 | 06-02-2022 | 2022 | 06-07-2022 |  |
| **a144** |  |  | INRAE | 47.83 | 1.91 | 06-02-2022 | 2022 | 06-03-2022 | yes |
| **a145** |  |  | INRAE | 47.83 | 1.91 | 06-02-2022 | 2022 | 06-07-2022 | yes |
| **a146** | *A. laticornis* | F | INRAE | 47.83 | 1.91 | 06-02-2022 | 2022 | 06-10-2022 |  |
| **a147** | *A. olivicolor* | F | University of Orléans | 47.85 | 1.94 | 06-02-2022 | 2022 | 06-07-2022 |  |
| **a148** | *A. laticornis* | F | INRAE | 47.83 | 1.91 | 06-03-2022 | 2022 | 06-13-2022 |  |
| **a149** | *A. laticornis* | M | INRAE | 47.83 | 1.91 | 06-03-2022 | 2022 | 06-09-2022 |  |
| **a150** | *A. angustulus* | F | University of Orléans | 47.85 | 1.94 | 06-03-2022 | 2022 | 06-08-2022 |  |
| **a151** | *A. laticornis* | M | University of Orléans | 47.85 | 1.94 | 06-07-2022 | 2022 | 06-09-2022 |  |
| **a152** | *A. laticornis* | F | University of Orléans | 47.85 | 1.94 | 06-07-2022 | 2022 | 06-13-2022 |  |
| **a153** | *A. laticornis* | M | University of Orléans | 47.85 | 1.94 | 06-07-2022 | 2022 | 06-10-2022 |  |
| **a154** | *M. fulgidicollis* | F | INRAE | 47.83 | 1.91 | 06-09-2022 | 2022 | 06-13-2022 |  |
| **a155** | *A. laticornis* | M | University of Orléans | 47.85 | 1.94 | 06-09-2022 | 2022 | 06-13-2022 |  |
| **a156** | *A. olivicolor* | F | University of Orléans | 47.85 | 1.94 | 06-10-2022 | 2022 | 06-15-2022 |  |
| **a157** | *A. laticornis* | M | INRAE | 47.83 | 1.91 | 06-10-2022 | 2022 | 06-14-2022 |  |
| **a158** | *A. laticornis* | F | INRAE | 47.83 | 1.91 | 06-13-2022 | 2022 | 06-16-2022 |  |
| **a159** | *A. laticornis* | F | University of Orléans | 47.85 | 1.94 | 06-13-2022 | 2022 | 06-17-2022 |  |
| **a160** | *A. laticornis* | M | INRAE | 47.83 | 1.91 | 06-14-2022 | 2022 | 06-16-2022 |  |
| **a161** | *A. laticornis* | M | INRAE | 47.83 | 1.91 | 06-14-2022 | 2022 | 06-20-2022 |  |
| **a162** |  |  | INRAE | 47.83 | 1.91 | 06-14-2022 | 2022 | 06-15-2022 |  |
| **a163** | *A. laticornis* | F | INRAE | 47.83 | 1.91 | 06-14-2022 | 2022 | 06-17-2022 |  |
| **a164** | *A. laticornis* | M | INRAE | 47.83 | 1.91 | 06-14-2022 | 2022 | 06-20-2022 |  |
| **a165** | *A. laticornis* | F | INRAE | 47.83 | 1.91 | 06-14-2022 | 2022 | 06-20-2022 |  |
| **a166** | *A. laticornis* | M | INRAE | 47.83 | 1.91 | 06-14-2022 | 2022 | 06-17-2022 |  |
| **a167** | *A. laticornis* | M | INRAE | 47.83 | 1.91 | 06-14-2022 | 2022 | 06-17-2022 |  |
| **a168** | *A. laticornis* | F | INRAE | 47.83 | 1.91 | 06-14-2022 | 2022 | 06-17-2022 |  |
| **a169** | *A. laticornis* | M | INRAE | 47.83 | 1.91 | 06-14-2022 | 2022 | 06-20-2022 |  |
| **a170** | *A. laticornis* | M | INRAE | 47.83 | 1.91 | 06-14-2022 | 2022 | 06-20-2022 |  |
| **a171** | *A. laticornis* | F | INRAE | 47.83 | 1.91 | 06-14-2022 | 2022 | 06-20-2022 |  |
| **a172** | *A. laticornis* | M | INRAE | 47.83 | 1.91 | 06-14-2022 | 2022 | 06-17-2022 |  |
| **a173** |  |  | INRAE | 47.83 | 1.91 | 06-14-2022 | 2022 | 06-15-2022 | yes |
| **a174** | *A. laticornis* | F | INRAE | 47.83 | 1.91 | 06-14-2022 | 2022 | 06-20-2022 |  |
| **a175** | *A. laticornis* | F | INRAE | 47.83 | 1.91 | 06-14-2022 | 2022 | 06-20-2022 |  |
| **a176** | *M. fulgidicollis* | M | INRAE | 47.83 | 1.91 | 06-14-2022 | 2022 | 06-16-2022 |  |
| **a177** | *A. laticornis* | M | INRAE | 47.83 | 1.91 | 06-14-2022 | 2022 | 06-17-2022 |  |
| **a178** | *A. laticornis* | F | INRAE | 47.83 | 1.91 | 06-14-2022 | 2022 | 06-16-2022 |  |
| **a179** | *A. laticornis* | F | INRAE | 47.83 | 1.91 | 06-14-2022 | 2022 | 06-17-2022 |  |
| **a180** | *A. laticornis* | M | INRAE | 47.83 | 1.91 | 06-14-2022 | 2022 | 06-17-2022 |  |
| **a181** | *A. laticornis* | M | University of Orléans | 47.85 | 1.94 | 06-14-2022 | 2022 | 06-16-2022 |  |
| **a182** | *A. laticornis* | F | University of Orléans | 47.85 | 1.94 | 06-14-2022 | 2022 | 06-20-2022 |  |
| **a183** | *A. olivicolor* | F | University of Orléans | 47.85 | 1.94 | 06-14-2022 | 2022 | 06-20-2022 |  |
| **a184** | *A. laticornis* | F | University of Orléans | 47.85 | 1.94 | 06-14-2022 | 2022 | 06-20-2022 |  |
| **a185** | *A. laticornis* | M | INRAE | 47.83 | 1.91 | 06-15-2022 | 2022 | 06-16-2022 |  |
| **a186** | *A. laticornis* | F | INRAE | 47.83 | 1.91 | 06-15-2022 | 2022 | 06-20-2022 |  |
| **a187** | *A. laticornis* | M | INRAE | 47.83 | 1.91 | 06-15-2022 | 2022 | 06-20-2022 |  |
| **a188** | *A. laticornis* | F | University of Orléans | 47.85 | 1.94 | 06-16-2022 | 2022 | 06-20-2022 |  |
| **a189** | *A. laticornis* | F | University of Orléans | 47.85 | 1.94 | 06-16-2022 | 2022 | 06-20-2022 |  |
| **a190** | *A. laticornis* | M | University of Orléans | 47.85 | 1.94 | 06-17-2022 | 2022 | 06-21-2022 |  |
| **a191** | *A. laticornis* | M | University of Orléans | 47.85 | 1.94 | 06-17-2022 | 2022 | 06-22-2022 |  |
| **a192** | *C. undatus* | F | University of Orléans | 47.85 | 1.94 | 06-20-2022 | 2022 | 06-23-2022 |  |
| **a193** | *A. laticornis* | F | University of Orléans | 47.85 | 1.94 | 06-21-2022 | 2022 | 06-23-2022 |  |
| **a194** |  |  | University of Orléans | 47.85 | 1.94 | 06-23-2022 | 2022 | 06-24-2022 | yes |
| **a195** | *A. olivicolor* | M | University of Orléans | 47.85 | 1.94 | 06-27-2022 | 2022 | 06-28-2022 |  |
| **a196** | *A. hyperici* | M | Ile Charlemagne | 47.90 | 1.97 | 06-26-2022 | 2022 | 07-05-2022 |  |
| **a197** | *A. hyperici* | M | Ile Charlemagne | 47.90 | 1.97 | 06-26-2022 | 2022 | 06-30-2022 |  |
| **a198** | *A. hyperici* | M | Ile Charlemagne | 47.90 | 1.97 | 06-26-2022 | 2022 | 06-28-2022 |  |
| **a199** | *A. hyperici* | F | Ile Charlemagne | 47.90 | 1.97 | 06-26-2022 | 2022 | 06-30-2022 |  |
| **a200** | *A. hyperici* | M | Ile Charlemagne | 47.90 | 1.97 | 06-26-2022 | 2022 | 06-30-2022 |  |
| **a201** | *A. hyperici* | F | Ile Charlemagne | 47.90 | 1.97 | 06-26-2022 | 2022 | 07-01-2022 |  |
| **a202** | *A. hyperici* | F | Ile Charlemagne | 47.90 | 1.97 | 06-26-2022 | 2022 | 06-30-2022 |  |
| **a203** | *A. hastulifer* | F | INRAE | 47.83 | 1.91 | 06-28-2022 | 2022 | 06-30-2022 |  |
| **a204** | *A. hyperici* | F | Ile Charlemagne | 47.90 | 1.97 | 06-28-2022 | 2022 | 07-01-2022 |  |
| **a205** | *A. hyperici* | M | Ile Charlemagne | 47.90 | 1.97 | 06-28-2022 | 2022 | 07-01-2022 |  |
| **a206** | *A. hyperici* | F | Ile Charlemagne | 47.90 | 1.97 | 06-28-2022 | 2022 | 07-01-2022 |  |
| **a207** | *A. hyperici* | F | Ile Charlemagne | 47.90 | 1.97 | 06-28-2022 | 2022 | 07-04-2022 |  |
| **a208** | *A. hyperici* | M | Ile Charlemagne | 47.90 | 1.97 | 06-28-2022 | 2022 | 07-04-2022 |  |
| **a209** | *A. hyperici* | M | Ile Charlemagne | 47.90 | 1.97 | 06-28-2022 | 2022 | 07-01-2022 |  |
| **a210** | *A. hyperici* | F | Ile Charlemagne | 47.90 | 1.97 | 06-28-2022 | 2022 | 07-04-2022 |  |
| **a211** | *A. hyperici* | F | Ile Charlemagne | 47.90 | 1.97 | 06-28-2022 | 2022 | 06-30-2022 |  |
| **a212** | *A. hyperici* | F | Ile Charlemagne | 47.90 | 1.97 | 06-28-2022 | 2022 | 07-05-2022 |  |
| **a213** | *A. hyperici* | M | Ile Charlemagne | 47.90 | 1.97 | 06-28-2022 | 2022 | 07-01-2022 |  |
| **a214** | *A. hyperici* | F | Ile Charlemagne | 47.90 | 1.97 | 06-28-2022 | 2022 | 07-01-2022 |  |
| **a215** | *A. hyperici* | F | Ile Charlemagne | 47.90 | 1.97 | 06-28-2022 | 2022 | 07-01-2022 |  |
| **a216** | *A. hyperici* | M | Ile Charlemagne | 47.90 | 1.97 | 06-28-2022 | 2022 | 07-04-2022 |  |
| **a217** | *A. hyperici* | F | Ile Charlemagne | 47.90 | 1.97 | 06-28-2022 | 2022 | 06-30-2022 |  |
| **a218** | *A. hyperici* | M | Ile Charlemagne | 47.90 | 1.97 | 06-28-2022 | 2022 | 07-04-2022 |  |
| **a219** | *A. hyperici* | M | Ile Charlemagne | 47.90 | 1.97 | 06-28-2022 | 2022 | 07-04-2022 |  |
| **a220** | *A. hastulifer* | F | INRAE | 47.83 | 1.91 | 06-29-2022 | 2022 | 07-04-2022 |  |
| **a221** | *A. graminis* | M | INRAE | 47.83 | 1.91 | 06-29-2022 | 2022 | 07-04-2022 |  |
| **a222** | *C. undatus* | F | INRAE | 47.83 | 1.91 | 06-29-2022 | 2022 | 07-04-2022 |  |
| **a223** | *A. laticornis* | F | University of Orléans | 47.85 | 1.94 | 06-30-2022 | 2022 | 07-04-2022 |  |
| **a224** | *A. laticornis* | F | INRAE | 47.83 | 1.91 | 07-01-2022 | 2022 | 07-04-2022 |  |
| **a225** | *A. laticornis* | F | University of Orléans | 47.85 | 1.94 | 07-01-2022 | 2022 | 07-05-2022 |  |
| **a226** | *A. laticornis* | F | University of Orléans | 47.85 | 1.94 | 07-04-2022 | 2022 | 07-06-2022 |  |
| **a227** | *A. laticornis* | F | University of Orléans | 47.85 | 1.94 | 07-04-2022 | 2022 | 07-07-2022 |  |
| **a228** | *A. hyperici* | M | University of Orléans | 47.85 | 1.94 | 07-04-2022 | 2022 | 07-07-2022 |  |
| **a229** | *A. hyperici* | M | University of Orléans | 47.85 | 1.94 | 07-04-2022 | 2022 | 07-15-2022 |  |
| **a230** | *A. hyperici* | F | University of Orléans | 47.85 | 1.94 | 07-04-2022 | 2022 | 07-05-2022 |  |
| **a231** | *A. laticornis* | F | University of Orléans | 47.85 | 1.94 | 07-06-2022 | 2022 | 07-11-2022 |  |
| **a232** | *A. hyperici* | F | INRAE | 47.83 | 1.91 | 07-06-2022 | 2022 | 07-12-2022 |  |
| **a233** | *A. graminis* | F | University of Orléans | 47.85 | 1.94 | 07-07-2022 | 2022 | 07-15-2022 |  |
| **a234** | *C. undatus* | M | University of Orléans | 47.85 | 1.94 | 07-07-2022 | 2022 | 07-11-2022 |  |
| **a235** | *A. laticornis* | M | University of Orléans | 47.85 | 1.94 | 07-07-2022 | 2022 | 07-11-2022 |  |
| **a236** | *A. laticornis* | M | University of Orléans | 47.85 | 1.94 | 07-07-2022 | 2022 | 07-11-2022 |  |
| **a237** | *A. graminis* | F | INRAE | 47.83 | 1.91 | 07-25-2022 | 2022 | 07-28-2022 |  |
| **a238** | *A. olivicolor* | F | INRAE | 47.83 | 1.91 | 07-25-2022 | 2022 | 07-27-2022 |  |
| **a239** | *A. graminis* | F | INRAE | 47.83 | 1.91 | 07-27-2022 | 2022 | 07-29-2022 |  |
| **a240** | *A. hyperici* | F | Vouzon field | 47.64 | 2.06 | 06-16-2023 | 2023 | 07-21-2023 |  |
| **a241** | *A. hyperici* | F | Vouzon field | 47.64 | 2.06 | 06-16-2023 | 2023 | 06-21-2023 |  |
| **a242** | *A. hyperici* | M | Vouzon field | 47.64 | 2.06 | 06-16-2023 | 2023 | 06-30-2023 |  |
| **a243** | *A. hyperici* | M | Vouzon field | 47.64 | 2.06 | 06-16-2023 | 2023 | 07-20-2023 |  |
| **a244** | *A. hyperici* | F | Vouzon field | 47.64 | 2.06 | 06-16-2023 | 2023 | 06-22-2023 |  |
| **a245** | *A. hyperici* | F | Vouzon field | 47.64 | 2.06 | 06-16-2023 | 2023 | 07-31-2023 |  |
| **a246** | *A. hyperici* | M | Vouzon field | 47.64 | 2.06 | 06-16-2023 | 2023 | 07-25-2023 |  |
| **a247** | *A. hyperici* | M | Vouzon field | 47.64 | 2.06 | 06-16-2023 | 2023 | 06-22-2023 |  |
| **a248** | *A. hyperici* | M | Vouzon field | 47.64 | 2.06 | 06-16-2023 | 2023 | 08-11-2023 |  |
| **a249** | *A. hyperici* | F | Vouzon field | 47.64 | 2.06 | 06-16-2023 | 2023 | 07-21-2023 |  |
| **a250** | *A. hyperici* | M | Vouzon field | 47.64 | 2.06 | 06-16-2023 | 2023 | 06-26-2023 |  |
| **a251** | *A. hyperici* | M | Vouzon field | 47.64 | 2.06 | 06-16-2023 | 2023 | 06-22-2023 |  |
| **a252** | *A. hyperici* | F | Vouzon field | 47.64 | 2.06 | 06-16-2023 | 2023 | 07-07-2023 |  |
| **a253** | *A. hyperici* | F | Vouzon field | 47.64 | 2.06 | 06-16-2023 | 2023 | 07-17-2023 |  |
| **a254** | *A. hyperici* | F | Vouzon field | 47.64 | 2.06 | 06-16-2023 | 2023 | 06-23-2023 |  |
| **a255** | *A. angustulus* | M | Vouzon field | 47.64 | 2.06 | 06-16-2023 | 2023 | 06-21-2023 |  |
| **a256** | *A. angustulus* | F | Vouzon forest | 47.65 | 2.01 | 06-16-2023 | 2023 | 06-20-2023 |  |
| **a257** | *A. angustulus* | F | Vouzon forest | 47.65 | 2.01 | 06-16-2023 | 2023 | 06-21-2023 |  |
| **a258** | *A. angustulus* | F | Vouzon forest | 47.65 | 2.01 | 06-16-2023 | 2023 | 06-20-2023 |  |
| **a259** | *A. angustulus* | F | Vouzon forest | 47.65 | 2.01 | 06-16-2023 | 2023 | 06-20-2023 |  |
| **a260** | *A. angustulus* | F | Vouzon forest | 47.65 | 2.01 | 06-16-2023 | 2023 | 06-21-2023 |  |
| **a261** | *A. angustulus* | M | Vouzon forest | 47.65 | 2.01 | 06-16-2023 | 2023 | 06-22-2023 |  |
| **a262** | *A. laticornis* | F | Vouzon forest | 47.65 | 2.01 | 06-16-2023 | 2023 | 06-22-2023 |  |
