## Supplementary material for "Intra- and interspecific variations in flight performance of oak-associated Agrilinae (Coleoptera: Buprestidae) using computerised flight mills": S2

Appendices

**Document S2.** Correlations among variables

Considering the large body size of *Coraebus undatus* compared to other species, global (i.e., non-species-specific) correlations among variables were analysed with and without *C. undatus* in the pool of species (Figure S2A). In both cases, strong correlations were found between distance variables (total flight distance, mean flight bout distance) and their duration counterparts (total flight duration, mean flight bout duration). Considering these strong correlations, duration variables were discarded from further analyses and emphasis was put on distance variables instead.

When *C. undatus* was included in the multidimensional clustering of variables, preflight mass clustered tightly with total flight distance, while the association became looser when excluding *C. undatus* from the pool. In addition, when *C. undatus* was excluded, preflight mass correlated with mean flight bout distance. The number of flight bouts was correlated with total flight duration or distance only when *C. undatus* was excluded, suggesting that other species performed multiple flights while *C. undatus* showed sustained flight bouts. With or without *C. undatus*, the latency to the first flight bout was inversely correlated with total flight distance.

**Figure S2A.** Correlation network among a selection of response variables measured for the first (eight-hour) trials, with all 12 species pooled together (left) and all but *C. undatus* (right). Highly correlated variables appear closer to each other, the proximity of points being determined by multidimensional clustering using a Principal Coordinates Analysis with *stats::cmdscale((1 - abs(correlations)), k = 2)* in *R* (see corrr package; Kuhn et al., 2022) and are joined by stronger paths. Path colour is determined by the correlation coefficient and sign, and correlations whose absolute value is below 0.2 are not shown.

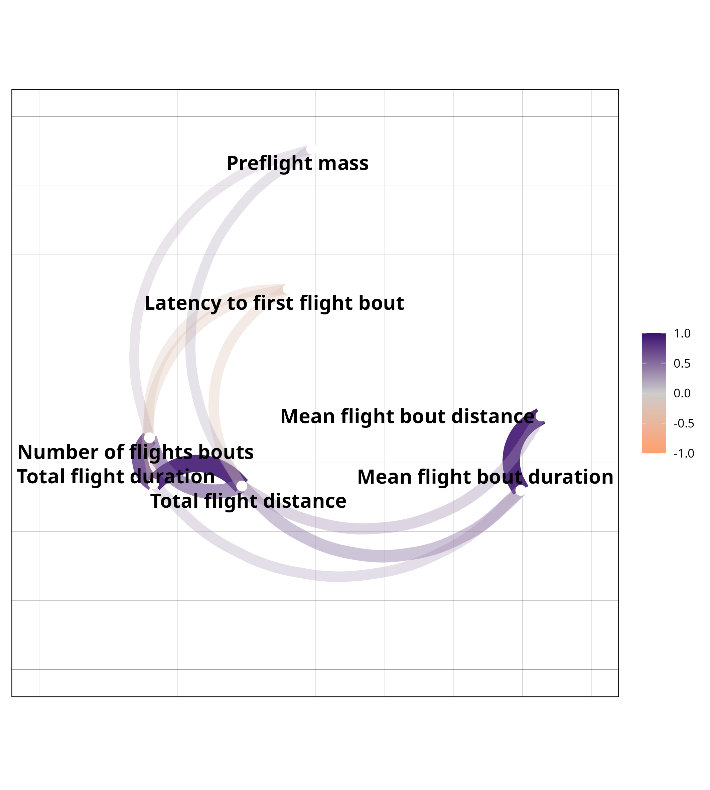

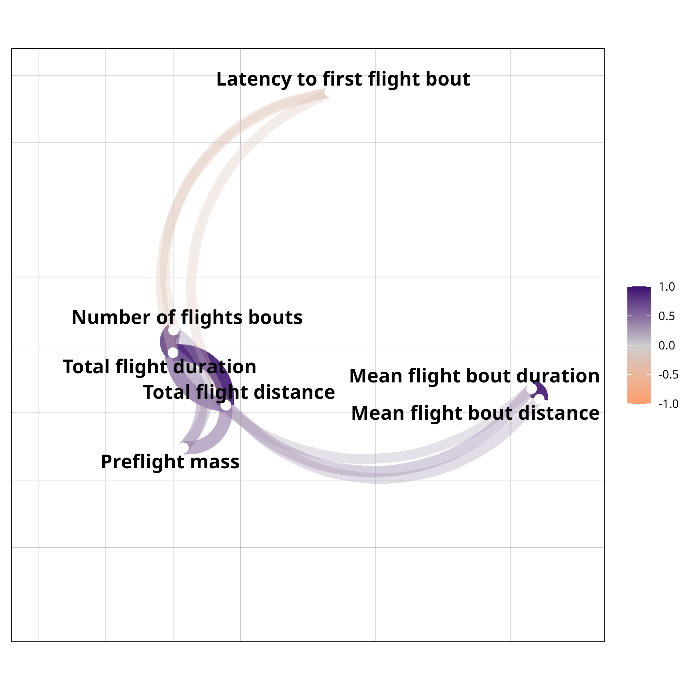

Given that species may show different flight strategies and are not equally represented in the sample, correlations with preflight mass were further analysed within the eight most numerous species only (Figure S2B). In all species but *Agrilus olivicolor*, higher preflight mass negatively impacted the latency to the first flight bout, in other words it increased the propensity to take off, or showed no relationship. This effect did not appear clearly in the multivariate analysis (Figure S2A), but could have been concealed by near-zero correlation coefficients in the two most abundant species, *A. laticornis* and *Agrilus hyperici*. Mixed effects with mean flight bout distance, total flight distance and number of flight bouts were found. Noticeable positive correlations were found for preflight mass with the former two variables in *C. undatus* only, and with the latter two variables in *Meliboeus fulgidicollis*, *Agrilus hyperici*, *A. laticornis* and *Agrilus hastulifer*. Some particularly high correlations coefficients found in *M. fulgidicollis* are, however, to be considered cautiously considering the low sample size and scarce active flights recorded in this species (four in total, see Figure S4), suggesting that such high coefficients resulted from a fraction of the individuals and could correspond to a biassed sample.

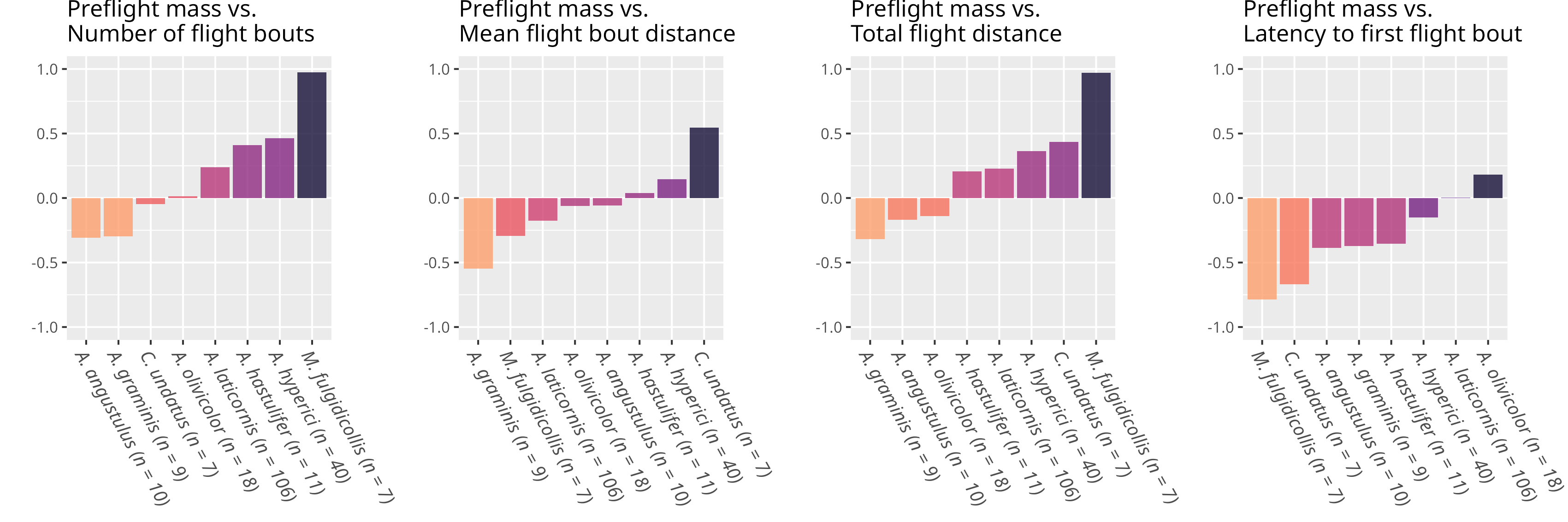

**Figure S2B.** Correlation coefficients between preflight mass and a selection of response variables during the first (eight-hour) trial, sorted by increasing value, in the eight most abundant species in the samples.
