## Supplementary material for "Intra- and interspecific variations in flight performance of oak-associated Agrilinae (Coleoptera: Buprestidae) using computerised flight mills": S3

Appendices

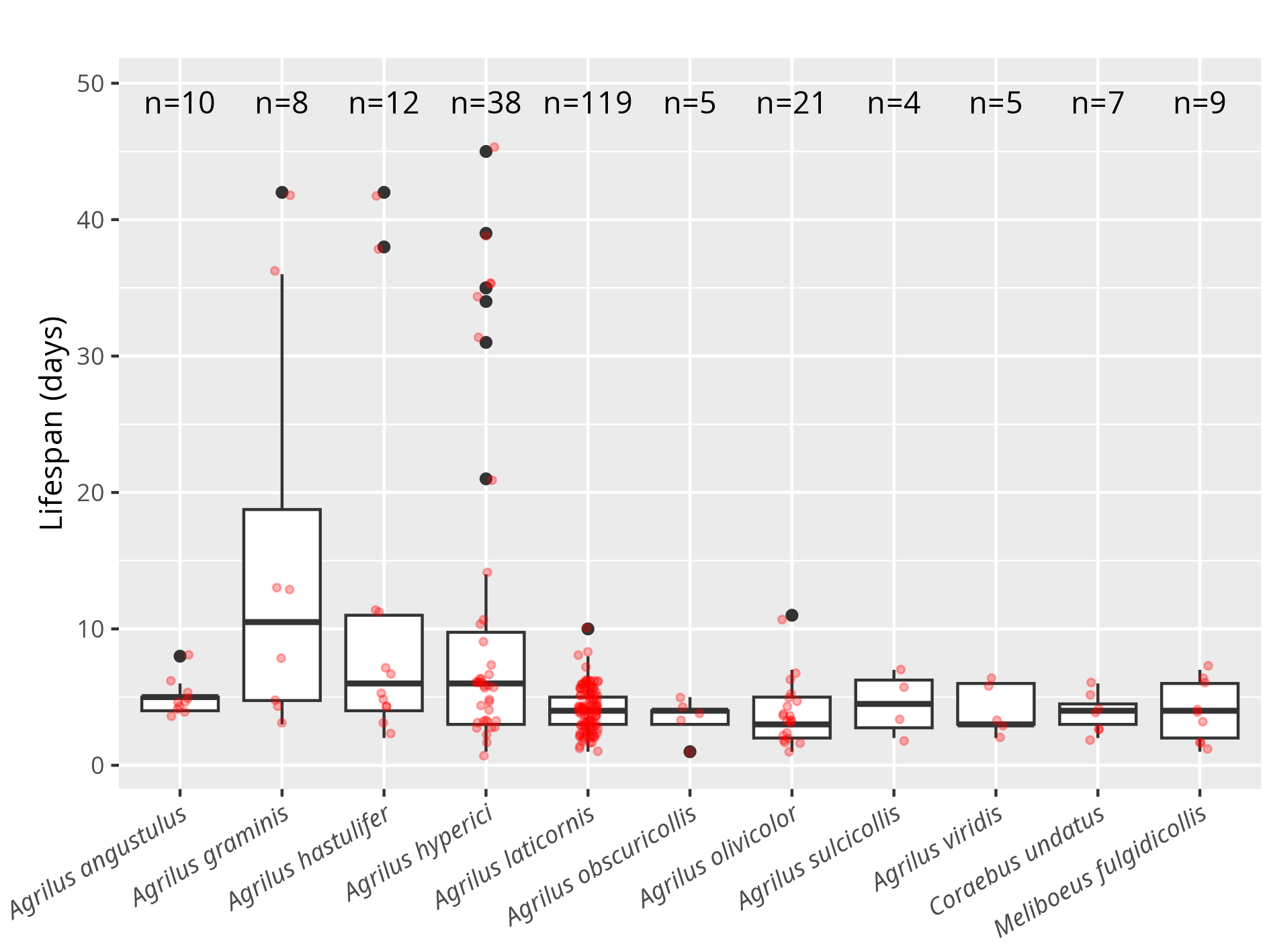

**Figure S3.** Lifespan of tested individuals in controlled conditions, after their capture. Lost or accidentally killed individuals were discarded. Red dots represent individual values, with random lateral offsets to improve visibility.
