## Supplementary material for "Intra- and interspecific variations in flight performance of oak-associated Agrilinae (Coleoptera: Buprestidae) using computerised flight mills": S4

Appendices

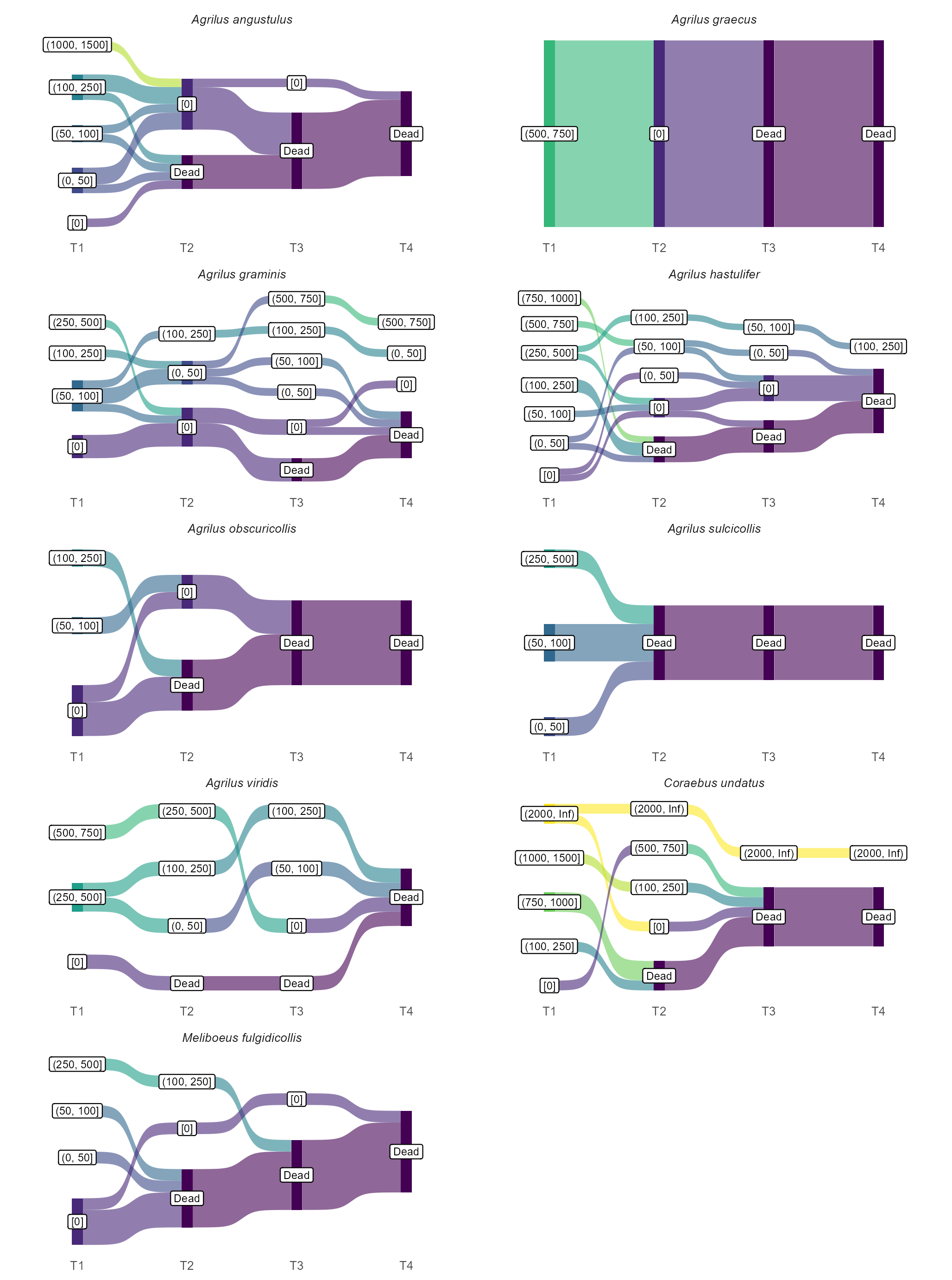

**Figure S4.** Individual flight trajectories across four consecutive trials in the less numerous species with the first trial spanning eight hours and the following four hours. Total flight distance is discretized in categories of increasing range to simplify individual trajectories.
